# Modification of the Neck Linker of KIF18A Alters Microtubule Subpopulation Preference

**DOI:** 10.1101/2023.05.02.539080

**Authors:** Katelyn A. Queen, Alisa Cario, Christopher L. Berger, Jason Stumpff

## Abstract

Kinesins support many diverse cellular processes, including facilitating cell division through mechanical regulation of the mitotic spindle. However, how kinesin activity is controlled to facilitate this process is not well understood. Interestingly, post-translational modifications have been identified within the enzymatic region of all 45 mammalian kinesins, but the significance of these modifications has gone largely unexplored. Given the critical role of the enzymatic region in facilitating nucleotide and microtubule binding, it may serve as a primary site for kinesin regulation. Consistent with this idea, a phosphomimetic mutation at S357 in the neck- linker of KIF18A alters the localization of KIF18A within the spindle from kinetochore microtubules to peripheral microtubules. Changes in localization of KIF18A-S357D are accompanied by defects in mitotic spindle positioning and the ability to promote mitotic progression. This altered localization pattern is mimicked by a shortened neck-linker mutant, suggesting that KIF18A-S357D may cause the motor to adopt a shortened neck-linker like state that prevents KIF18A from accumulating at the plus-ends of kinetochore microtubules. These findings demonstrate that post-translational modifications in the enzymatic region of kinesins could be important for biasing their localization to particular microtubule subpopulations.

## INTRODUCTION

Kinesins were first identified as proteins that transport organelles via microtubules in squid giant axons (Vale et al., 1985). Since their initial identification, extensive work has highlighted the critical role kinesins play in several cellular functions ranging from intracellular transport (Hirokawa et al., 2009), to cell division (Vicente and Wordeman, 2015; Cross and McAinsh, 2014), and cell migration (Hirokawa et al., 2009; Bachmann and Straube, 2015). These functions are carried out in mammalian organisms by forty-five kinesins, which share a conserved structure consisting of a force producing motor domain, a conformational-shifting neck-linker, a coiled-coil region that promotes dimerization, and a tail which facilitates kinesin function by interacting with cargo or microtubules (Hirokawa, 1998; Verhey and Hammond, 2009). How the conserved structures of kinesins are regulated for a diverse array of cellular functions in both a temporal and spatial-dependent manner remains incompletely understood.

Several mechanisms of regulation have been identified that work to control the function of individual kinesins. For example, autoregulation, whereby conformational shifts in the kinesin structure result in interactions between the kinesin motor domain and tail, prevent enzymatic activation (Coy et al., 1999; Verhey et al., 1998; Hirokawa et al., 1989). This autoinhibition is subsequently released by cargo binding to the kinesin tail and changes in ionic strength (Hackney et al., 1992; Verhey et al., 1998; Coy et al., 1999). Other mechanisms of regulation include preferential interactions with specific microtubule isotypes and post-translational modifications (PTMs) (Nakata and Hirokawa, 2003; Sirajuddin et al., 2014; Reed et al., 2006; Kaul et al., 2014), cell-cycle-dependent expression (Mayr et al., 2007; Brown et al., 1994; Funabiki and Murray, 2000), and inhibition by kinesin binding protein (Kevenaar et al., 2016; Malaby et al., 2019b; Atherton et al., 2020; Solon et al., 2021).

Many proteins are known to be regulated through PTMs, which can reversibly control their function in a temporal and spatial dependent manner (Nishi et al., 2014; Olsen et al., 2006; Deribe et al., 2010). Large-scale mass spectrometry studies have identified post-translational modifications within the conserved enzymatic region of all forty-five mammalian kinesins (Hornbeck et al., 2015). However, the functional relevance of the majority of these modifications remains unknown. Studies that have examined motor domain modifications indicate that they do indeed alter kinesin function. For example, acetylation of the KIF11 alpha-2 helix causes the motor to resist microtubule sliding (Muretta et al., 2018), phosphorylation of the KIF11 alpha-3 helix results in reduced mitotic activity (Bickel et al., 2017), and phosphorylation of the kinesin-1 alpha-3 helix tunes the directionality of cargo by decreasing stall force and velocity (DeBerg et al., 2013).

Modifications have been also reported within the neck-linkers of several kinesins including KIF11, KIF23, KIF20A, KIF26A, KIF3A, KIF2A, and KIF18A (Hornbeck et al., 2015). This structurally conserved region is critically important for processive movement and force generation by kinesins. In addition, neck-linker length directly affects the ability of kinesin motors to navigate obstacles on the microtubule (Malaby et al., 2019a; Shastry and Hancock, 2010; Hoeprich et al., 2014). Thus, modifications of the neck linker could change the mechanical output or agility of a kinesin motor.

KIF18A is a mammalian mitotic kinesin that accumulates on the plus-ends of kinetochore microtubules, where it suppresses microtubule dynamics to promote chromosome alignment and mitotic progression (Du et al., 2010; Gardner et al., 2008; Stumpff et al., 2008; Mayr et al., 2007; Stumpff et al., 2012; Janssen et al., 2018). Additionally, KIF18A is required to maintain spindle length in mitosis (Weaver et al., 2011; Goshima et al., 2005) and accumulates on bridging fibers where it is thought to play a role in regulating bridging fiber dynamics (Jagrić et al., 2021). While KIF18A function has been primarily studied in mitosis, KIF18A nuclear localization sequence mutants have been shown to decrease the dynamicity of microtubule ends in interphase cells (Du et al., 2010), and loss of KIF18A affects cell migration (Zhang et al., 2010; Qian et al., 2021) and the control of microtubule dynamics in neurons (Kevenaar et al., 2016). How KIF18A’s activity is regulated for these various roles is not understood.

Serine 357 (S357) within the neck-linker of KIF18A was identified as a phosphorylation site via mass spectrometry (Imami et al., 2008). We show here that an S357D mutation alters the localization of KIF18A, displacing the motor from kinetochore microtubules to peripheral microtubules of the mitotic spindle. This relocalization causes spindle rotation but only has a moderate effect on the canonical mitotic functions of KIF18A. We demonstrate that the altered accumulation of the motor to peripheral microtubules does not result in changes to microtubule dynamics, microtubule density, or the velocity at which the motor moves along microtubules. Interestingly, the localization pattern of the S357D mutant closely mimics that of a shortened neck-linker (sNL) mutant that also reduces KIF18A accumulation at the plus-ends of kinetochore microtubules (Malaby et al., 2019a). Removal of the kinetochore microtubule bundling protein HURP allows for some localization of both the sNL and S357D mutants to kinetochore microtubules but does not fully rescue localization away from peripheral microtubules. We hypothesize that charge at S357 essentially kinks the neck-linker of KIF18A, creating a shortened neck-linker like state, which results in altered accumulation of KIF18A to the peripheral microtubules. These findings demonstrate how a single modification in the neck-linker can alter KIF18A localization and potentially serve as a mechanism for directing the motor to a specific subset of microtubules.

## RESULTS

### KIF18A S357D results in altered localization

Serine 357 of KIF18A is conserved across kinesin-8 homologs and was previously identified as a site of phosphorylation (Figure 1a) (Imami et al., 2008). To study how phosphorylation of S357 might affect KIF18A activity and function, we generated HeLa Kyoto and RPE1 cell lines that express inducible, siRNA-resistant, and GFP-tagged wild-type, S357A (phosphonull), or S357D (phosphomimetic) KIF18A. Given that previous work has highlighted differential dependence on KIF18A for proliferation (Marquis et al., 2021; Cohen-Sharir et al., 2021; Quinton et al., 2021) both HeLa Kyoto and RPE1 cell lines were generated to compare phenotypes in aneuploid cancer cells to near diploid cells. KIF18A protein expression levels were found to be comparable across the different cell lines (Supplementary Figure S1). To examine localization of S357 mutants, cells were treated with siRNA to deplete endogenous KIF18A and doxycycline to induce expression of GFP-tagged KIF18A variants and then fixed 24 hours later. Wild-type GFP-KIF18A, in both HeLa Kyoto and RPE1 cells, displayed a canonical localization pattern with accumulation at the plus-ends of kinetochore microtubules in metaphase cells. GFP-KIF18A S357A displayed a similar localization pattern. However, GFP-KIF18A S357D accumulated on peripheral microtubules in both HeLa Kyoto and RPE1 cells (Figure 1b). This altered accumulation was more drastic in RPE1 cells, where most of the kinetochore microtubule localization was lost and KIF18A S357D was observed primarily on the peripheral spindle microtubules (Figure 1b).

**Figure 1:**
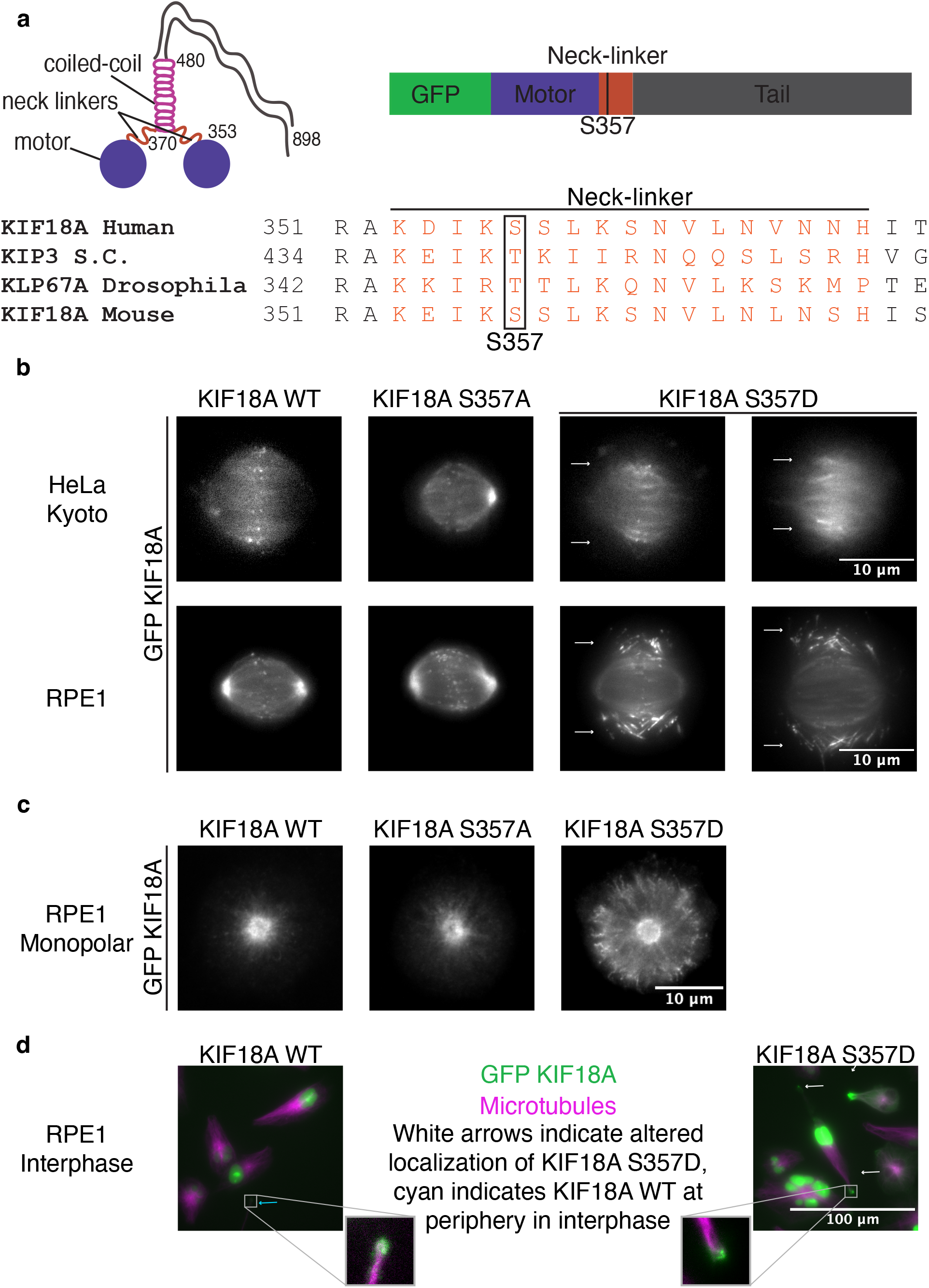
KIF18A S357D exhibits altered localization. **(a)** Cartoon of KIF18A dimer with important structural regions noted and sequence alignment of KIF18A homologs indicating conservation of S357 site. S.C.: Saccharomyces cerevisiae. **(b)** Representative immunofluorescence images of GFP KIF18A wild-type (WT), S357A, and S357D localization in bipolar HeLa Kyoto and RPE1 cells. Cells were fixed approximately 24 hours after knockdown of endogenous KIF18A and induction of GFP- KIF18A with doxycycline. White arrows indicate altered localization of KIF18A S357D. Brightness/contrast levels set differently to optimize visualization. Scale bar is 10 μm. **(c)** Representative immunofluorescence images of GFP-KIF18A localization in monopolar mitotic cells. Cells were fixed approximately 24 hours after siRNA treatment to knockdown endogenous KIF18A and induction of GFP-KIF18A with doxycycline. 2-3 hours prior to fixation, monopolar mitotic cells were established with addition of 100 μm monastrol. Brightness/contrast levels set differently to optimize visualization. WT: wild-type. Scale bar is 10 μm. **(d)** Representative immunofluorescence images of KIF18A wild-type (WT) and S357D localization in interphase cells. Cells were imaged live approximately 24 hours after siRNA treatment to knockdown endogenous KIF18A and induction of GFP-KIF18A with doxycycline. Colors indicate pseudo-color in merged image. Brightness/contrast levels set differently to optimize visualization. Cyan arrows indicate WT KIF18A localization at periphery of interphase cells, white arrows indicate additional accumulation of KIF18A S357D at periphery. Insets created with increased brightness and contrast to highlight peripheral localization. Scale bar is 100μm. WT: wild-type.

Treating cells with low doses of the microtubule stabilizing drug taxol results in enhanced accumulation of wild-type KIF18A to the plus-ends of kinetochore microtubules (Stumpff et al., 2011). This additional accumulation of KIF18A is likely due to decreases in microtubules dynamics, which allow additional time for KIF18A accumulation at plus-ends. To test if the altered localization of the KIF18A S357D mutant could be rescued when microtubules were further stabilized, we treated cells with low doses of taxol and imaged KIF18A localization over time. Results indicate that taxol-stabilization of the microtubules did not rescue the altered accumulation of KIF18A S357D on peripheral microtubules (Supplementary Figure S2 and Videos 1-3). This work suggests that the altered accumulation of KIF18A S357D is independent of microtubule dynamics.

To further examine the altered localization of KIF18A S357D, we analyzed KIF18A localization in both monopolar mitotic cells and interphase cells. Wild-type and S357A KIF18A were primarily localized near the centrosomes of monopolar spindles while KIF18A S357D accumulated at the peripheral edge of monopolar spindles (Figure 1c). In interphase cells, wild- type KIF18A was primarily nuclear due to a nuclear localization sequence. However, a small population of wild-type KIF18A was observed at the peripheral edge of some interphase cells (Figure 1d). Interestingly, cells expressing the KIF18A S357D mutant also displayed an increase in the population of KIF18A observed at the peripheral edge of interphase cells (Figure 1d). This work indicates that the S357D mutation alters the microtubules KIF18A accumulates on in both mitosis and interphase and that this altered localization is not dependent on formation of a bipolar mitotic spindle.

### KIF18A S357D causes spindle rotation

Proper cell division requires the mitotic spindle to maintain a centralized position within the dividing cell. The stabilization of the mitotic spindle depends on contact of peripheral microtubules with the cortex (Garzon-Coral et al., 2016; Grill and Hyman, 2005), cortical dynein forces centering microtubule asters (Laan et al., 2012), and actin and myosin forces acting on the mitotic cell (Fink et al., 2011). Thus, we investigated whether the altered accumulation of KIF18A- S357D to peripheral microtubules impacted the stabilization of the mitotic spindle in dividing cells. In cells expressing KIF18A S357D, we observed rotation of the mitotic spindle (Figure 2a, 2b and Videos 4-5). These spindle rotations occurred in live-cells in the presence or absence of siR-tubulin labeled microtubules (Videos 6-7). While there was an increase in spindle rotation in the presence of KIF18A S357D, no change in the spindle angle relative to the focal plane was observed in fixed cells (Figure 2c), indicating that the spindle rotation occurs primarily in the xy-plane. This movement of the mitotic spindle suggests that the altered accumulation of KIF18A S357D impacts the balance of forces necessary to maintain a stabilized spindle position during cell division.

**Figure 2:**
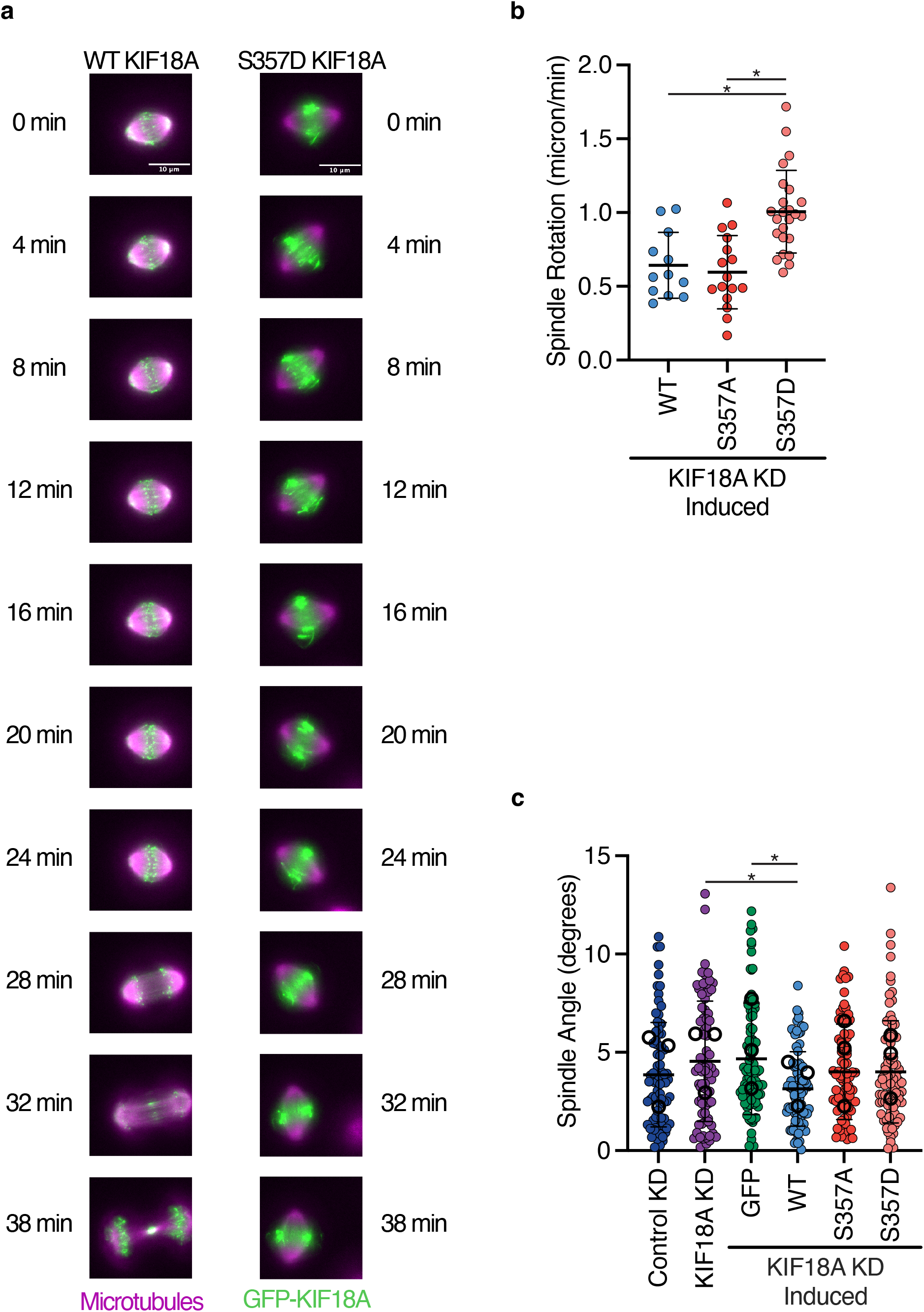
KIF18A S357D causes defects in mitotic spindle centering. **(a)** Representative still images from live-cell imaging of GFP-KIF18A wild-type (WT) and S357D. Cells were imaged approximately 24 hours after siRNA treatment and induction of GFP-KIF18A with doxycycline and 3 hours after addition of siR-tubulin to label microtubules. Colors indicate pseudo-color in merged images. Brightness/contrast levels are set to be equivalent. Scale bar is 10 μm. **(b)** Quantification of spindle rotation in cells expressing wild-type (WT), S357A, or S357D KIF18A. Solid horizontal line indicates mean, vertical lines indicate standard deviation. Each dot represents a single cell. Data were acquired from three experimental replicates. Data were analyzed using a one-way ANOVA with Tukey’s test for multiple comparisons. P value style: <0.05 (*), if no significance is indicated result was not significant ( > 0.05). KD: knockdown. **(c)** Quantification of the spindle angle calculated from the dot product between the two vectors resulting from the x, y, z coordinates of each spindle pole. Solid horizontal line indicates mean, vertical lines indicate standard deviation. Each small solid dot represents a single cell, larger open dots represent averages from each experimental replicate. Data were acquired from three experiments. Data were analyzed using a one-way ANOVA with Tukey’s test for multiple comparisons. P value style: <0.05 (*), if no significance is indicated result was not significant ( > 0.05). KD: knockdown.

### KIF18A S357D reduces the ability of KIF18A to promote chromosome alignment and maintain spindle length

KIF18A activity is required for chromosome alignment and maintaining spindle length (Du et al., 2010; Gardner et al., 2008; Stumpff et al., 2008, 2012; Mayr et al., 2007). To examine how the altered accumulation of KIF18A S357D impacts the canonical mitotic functions of KIF18A, we treated cells with siRNAs targeting KIF18A and induced expression of wild-type, S357A, or S357D KIF18A. To measure chromosome alignment, the distribution of kinetochore intensity along the spindle pole axis was fit to a gaussian distribution. The width of the gaussian distribution at half the maximum kinetochore fluorescence intensity was then determined (Fonseca and Stumpff, 2016; Malaby et al., 2019b). Both wild-type and S357A KIF18A were able to promote chromosome alignment following depletion of endogenous KIF18A in both HeLa Kyoto and RPE1 cells (Figure 3a-b and S3a-b). Interestingly, while KIF18A S357D was less efficient than wild-type KIF18A at promoting chromosome alignment, the mutant motor was able to facilitate more extensive chromosome alignment compared to KIF18A knockdown (Figure 3a-b and S3a-b).

**Figure 3:**
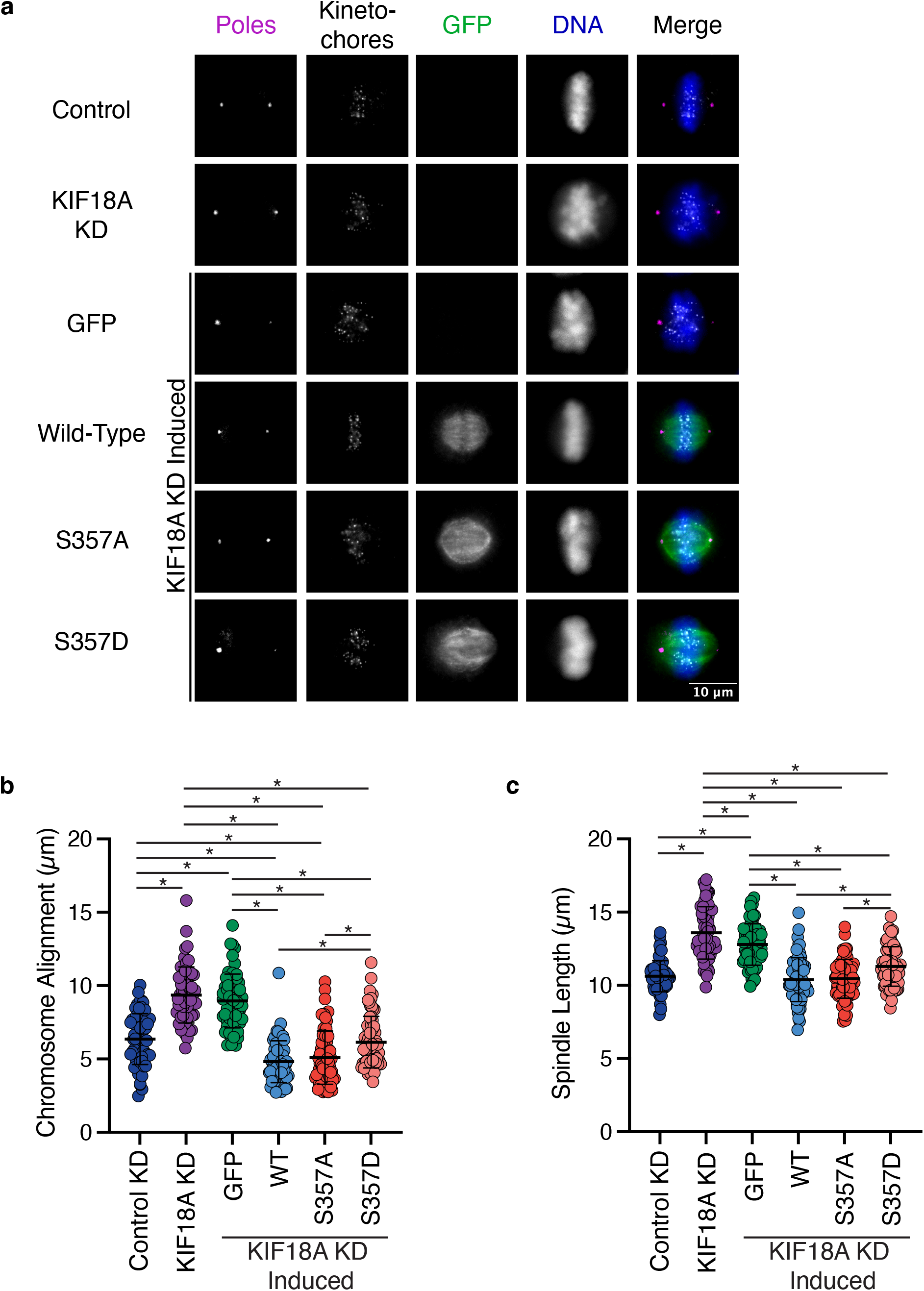
KIF18A S357D displays reduced chromosome alignment and spindle length control in HeLa Kyoto cells. **(a)** Representative immunofluorescence images of HeLa Kyoto cells. Cells were fixed approximately 24 hours after siRNA treatment to knockdown endogenous KIF18A and induction of GFP-KIF18A with doxycycline. Color indicates pseudo-color in merged image. Brightness/contrast levels for GFP are set to be equivalent, brightness/contrast levels for poles, kinetochore, and DNA are set differently to optimize visualization. Scale bar is 10 μm. KD: knockdown. **(b)** Quantification of chromosome alignment in HeLa Kyoto cells. Chromosome alignment was determined by measuring the distribution of kinetochores between spindle poles in a metaphase cell. This distribution was fit to a gaussian curve, and the width of the distribution at half the maximum fluorescence intensity was recorded as the value for chromosome alignment. Solid horizontal line indicates mean, vertical lines indicate standard deviation. Each dot represents a single cell. Data were acquired from three experimental replicates. Data were analyzed via a one-way ANOVA with Tukey’s test for multiple comparisons. P value style: <0.05 (*), if no significance is indicated result was not significant (> 0.05). KD: knockdown, WT: wild-type. **(c)** Quantification of spindle length in HeLa Kyoto cells. Solid horizontal line indicates mean, vertical lines indicate standard deviation. Each dot represents a single cell. Data were acquired from three experimental replicates. Data were analyzed via a one-way ANOVA with Tukey’s test for multiple comparisons. P value style: <0.05 (*), if no significance is indicated result was not significant (> 0.05). KD: knockdown, WT: wild-type.

Similar effects were observed with respect to spindle length control, which was measured by determining the distance between centrosomes in bipolar metaphase cells. Both wild-type and S357A KIF18A were able to promote spindle length maintenance (Figure 3c and S3c). In contrast, KIF18A S357D was not as efficient as wild-type KIF18A at maintaining spindle length but was able to reduce spindle length compared to spindles in KIF18A knockdown cells (Figure 3c and S3c). These results indicate that while the mitotic functions of KIF18A S357D are compromised compared to wild-type KIF18A, the mutant is partially functional.

### KIF18A S357D is not competent to promote mitotic progression

In addition to causing an increase in unaligned chromosomes, depletion of KIF18A leads to an increase in the time needed to complete mitosis and mitotic arrest in some cell types (Janssen et al., 2018; Czechanski et al., 2015; Marquis et al., 2021; Fonseca et al., 2019). To determine how the altered localization of KIF18A S357D impacts the ability of KIF18A to promote mitotic progression, we performed a series of live-cell imaging experiments and quantified the average time cells took to go from nuclear envelope breakdown to anaphase onset (Figure 4a), as well as the percent of cells that arrested in mitosis (entered mitosis but never divided) in both HeLa Kyoto and RPE1 cells. As observed in previous studies (Stumpff et al., 2008; Mayr et al., 2007; Zhu et al., 2005), knockdown of KIF18A in HeLa Kyoto cells results in an increase in the time it takes for cells to proceed through mitosis (Figure 4b). This increase in mitotic timing was rescued in cells expressing either wild-type or S357A KIF18A (Figure 4b). However, KIF18A S357D was unable to promote mitotic progression, leading to an increase in mitotic timing comparable to that measured following KIF18A knockdown (Figure 4b). In contrast to the extended mitotic delay in HeLa cells, as has been shown previously (Fonseca et al., 2019), KIF18A knockdown in RPE1 results in a modest increase in mitotic timing (Figure 4c). Despite this, KIF18A S357D expression led to a slight increase in mitotic timing compared to timing in control cells or those expressing wild-type and or S357A KIF18A following KIF18A knockdown (Figure 4c).

**Figure 4:**
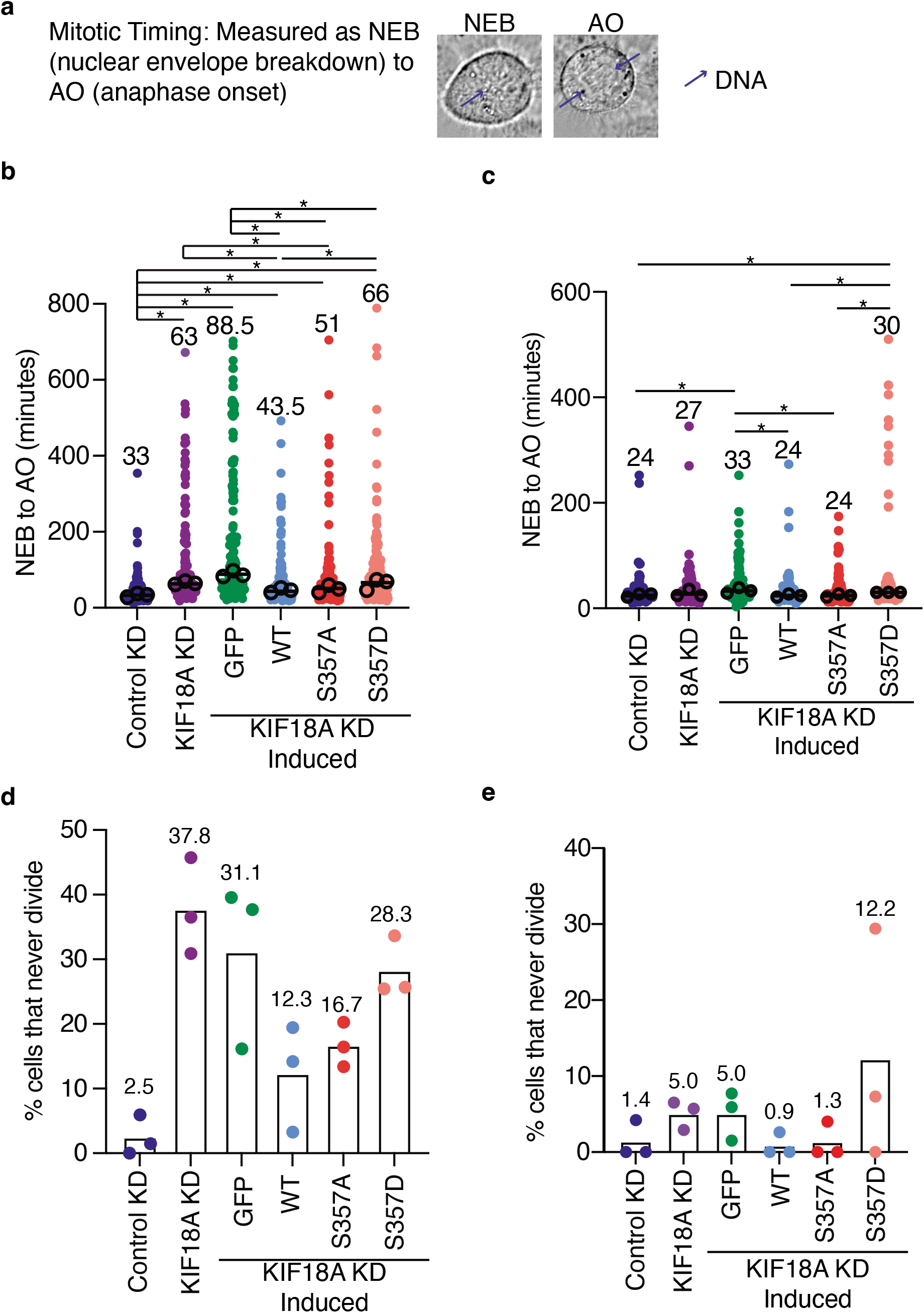
KIF18A S357D causes defects in mitotic progression. **(a)** Representative DIC images highlighting quantification of nuclear envelope breakdown (NEB) to anaphase onset (AO). Blue arrows indicate DNA location in cell. **(b-c)** Quantification of NEB to AO (mitotic timing) in HeLa Kyoto cells **(b)** and RPE1 cells **(c)**. Horizontal line represents median. Each solid small dot represents a single cell, larger open circles represent the average from each experimental replicate. Data were acquired from three experimental replicates and were analyzed using a Kruskal-Wallis with Dunn’s Multiple Comparisons Test. P value style: P value style: <0.05 (*), if no significance is indicated result was not significant (> 0.05). Numbers above each condition indicate the median mitotic timing value. WT: wild-type, KD: knockdown. **(d-e)** Quantification of the percent of cells that never divide in HeLa Kyoto cells **(d)** and RPE1 cells **(e)**. Bar indicates mean, each dot represents the percentage of cells that never divided for an individual experimental replicate. Numbers above graphs indicate the mean overall value for each condition. WT: wild-type, KD: knockdown.

KIF18A knockdown in HeLa Kyoto cells results in a population of cells that arrest in mitosis (37.8%, Figure 4d). This fraction of arrested cells is partially rescued by expression of wild-type or S357A KIF18A, while KIF18A S357D is less efficient at promoting mitotic progression (Figure 4d). In RPE1 cells, KIF18A knockdown results in a modest increase in mitotic arrest (Figure 4e), which is consistently rescued upon the induction of wild-type or S357A KIF18A. However, induction of KIF18A S357D failed to reduce the population of arrested RPE1 cells (Figure 4e). These data indicate that while KIF18A S357D retains partial chromosome alignment and spindle length maintenance functions, it is unable to promote mitotic progression.

### KIF18A S357D does not affect spindle microtubule organization or motor velocity

Kinetochore microtubules are relatively stable compared to non-kinetochore microtubules within the spindle (Girão and Maiato, 2019; Karsenti and Vernos, 2001). The increased stability of kinetochore microtubules is believed to contribute to the accumulation of wild-type KIF18A at the plus ends of this subset of microtubules (Stumpff et al., 2011). It is possible that the altered localization of the KIF18A S357D mutant stabilizes the peripheral microtubules resulting in the observed localization phenotype. If this were the case, we would expect there to be a decrease in the dynamics of peripheral microtubules in cells expressing the KIF18A S357D mutant compared to the wild-type motor. To test if KIF18A S357D expression induced changes in the dynamics of spindle or peripheral microtubules, we performed a series of immunofluorescence experiments to quantify the density and polymerization levels of spindle and peripheral microtubules. To analyze the fraction of polymerizing microtubules, as a proxy for microtubule dynamics, RPE1 cells were stained for the end-binding protein EB1. EB1 is a microtubule tip tracking protein that associates with the growing tip of a microtubule (Nehlig et al., 2017). If KIF18A S357D reduced the dynamics of peripheral microtubules, we would expect a subsequent reduction in the population of EB1-labeled microtubules at the periphery of the spindle. To simplify quantification, these experiments were conducted in monopolar cells, where altered KIF18A S357D localization is still observed. Results indicate that despite KIF18A S357D localization to the peripheral edge of a monopolar spindle, the population of polymerizing, EB1- labeled microtubule ends are similar in WT KIF18A and KIF18A S357D expressing cells (Figure 5a- c).

**Figure 5:**
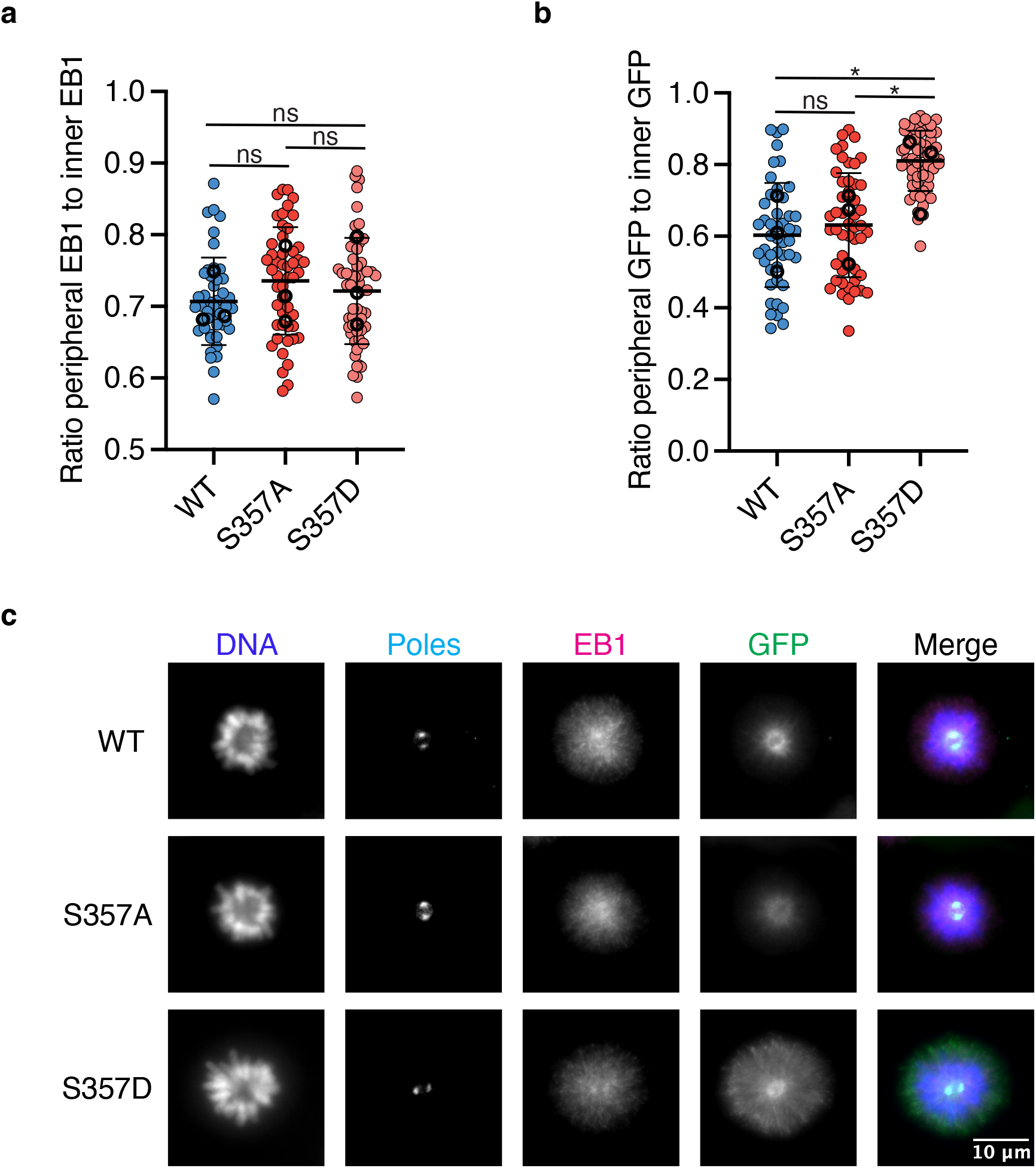
KIF18A S357D does not alter peripheral microtubule polymerization. **(a)** Quantification of the ratio of peripheral EB1 signal to inner (spindle) EB1 signal. Solid horizontal line indicates mean, vertical lines indicate standard deviation. Each solid small dot represents a single cell, larger open dots represent the mean from each experimental replicate. Data were acquired from three experimental replicates and were analyzed using a one-way ANOVA with Tukey’s test for multiple comparisons. P value style: >0.05 (ns), if <0.05 (*). WT: wild-type. **(b)** Quantification of the ratio of peripheral GFP signal to inner (spindle) GFP signal. Solid horizontal line indicates mean, vertical lines indicate standard deviation. Each solid small dot represents a single cell, larger open dots represent the mean from each experimental replicate. Data were acquired from three experimental replicates and were analyzed using a one-way ANOVA with Tukey’s test for multiple comparisons. P value style: >0.05 (ns), if <0.05 (*). WT: wild-type. **(c)** Representative immunofluorescence images of EB1 and GFP in RPE1 cells. Cells were fixed approximately 24 hours after siRNA treatment to knockdown endogenous KIF18A and induction of GFP-KIF18A with doxycycline. 2-3 hours prior to fixation monopolar mitotic cells were established with addition of 100 μm monastrol. Color indicates pseudo-color in merged image. Brightness/contrast levels for GFP and EB1 are set to be equivalent, brightness/contrast levels for poles and DNA are set differently to optimize visualization. Scale bar is 10 μm.

If the population of peripheral microtubules that the KIF18A S357D mutant localizes to were stabilized by the motor, we would expect to see an increase in the density of peripheral microtubules in KIF18A S357D expressing cells. To test this, we fixed and stained RPE1 cells for tubulin and measured the tubulin fluorescence at the spindle periphery and within the main body of the spindle. The resulting data indicate that tubulin immunofluorescence was similar within the spindles and spindle peripheries of cells expressing wild type KIF18A or KIF18A S357D (Figure 6a-d). However, we did observe a reduction in tubulin immunofluorescence with the KIF18A S357A mutant (Figure 6c-d). Taken together with the EB1 analyses, these data indicate that the altered accumulation of KIF18A S357D does not accompany gross changes in microtubule polymerization or number. This would also suggest that the peripheral microtubules that KIF18A S357D localizes to are present in control cells.

**Figure 6:**
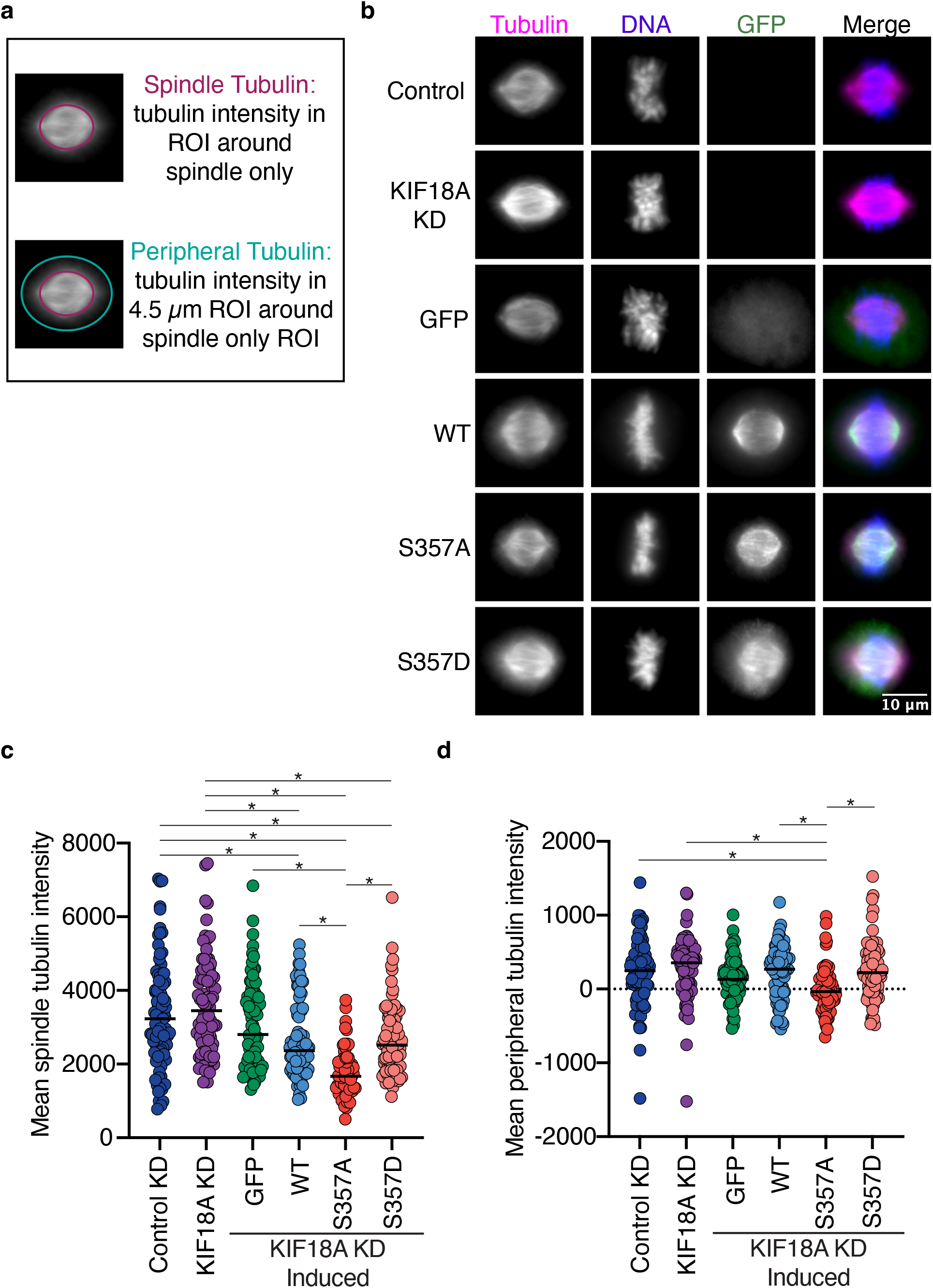
KIF18A S357D does not change microtubule density. **(a)** Graphic demonstrating quantification of spindle and peripheral tubulin. **(b)** Representative images of tubulin, DNA, and GFP in RPE1 cells. Cells were fixed approximately 24 hours after siRNA treatment to knockdown endogenous KIF18A and induction of GFP-KIF18A with doxycycline. Color indicates pseudo-color in merged image. Brightness/contrast levels for tubulin are set to be equivalent, brightness/contrast levels for DNA and GFP are set differently to optimize visualization. Scale bar is 10 μm. WT: wild-type, KD: knockdown. **(c-d)** Quantification of background subtracted spindle tubulin **(c)** and peripheral tubulin **(d)** intensity in RPE1 cells. Horizontal line represents median. Each dot represents a single cell. Data were acquired from a minimum of three experimental replicates and were analyzed using a Kruskal-Wallis with Dunn’s Multiple Comparisons Test. P value style: <0.05 (*), if no significance is indicated result was not significant (> 0.05). WT: wild- type, KD: knockdown.

Another possible explanation for the altered localization of KIF18A S357D is that increased motor velocity could allow it to reach the ends of faster polymerizing, non-kinetochore microtubules. To test this hypothesis, we performed single molecule total internal reflection fluorescence microscopy (TIRF) experiments to quantify the velocity of KIF18A wild-type and S357D motors on both taxol stabilized and dynamic microtubules. Results indicate that on both stabilized and dynamic microtubules KIF18A S357D moves at a similar velocity to wild type KIF18A (Figure 7a-d). Surprisingly, both KIF18A wild-type and KIF18A S357D displayed an increased velocity on dynamic microtubules compared to taxol stabilized microtubules (Figure 7). This work suggests that the altered localization of KIF18A S357D is not due to the motor moving at a faster velocity, allowing for accumulation on the peripheral microtubules.

**Figure 7:**
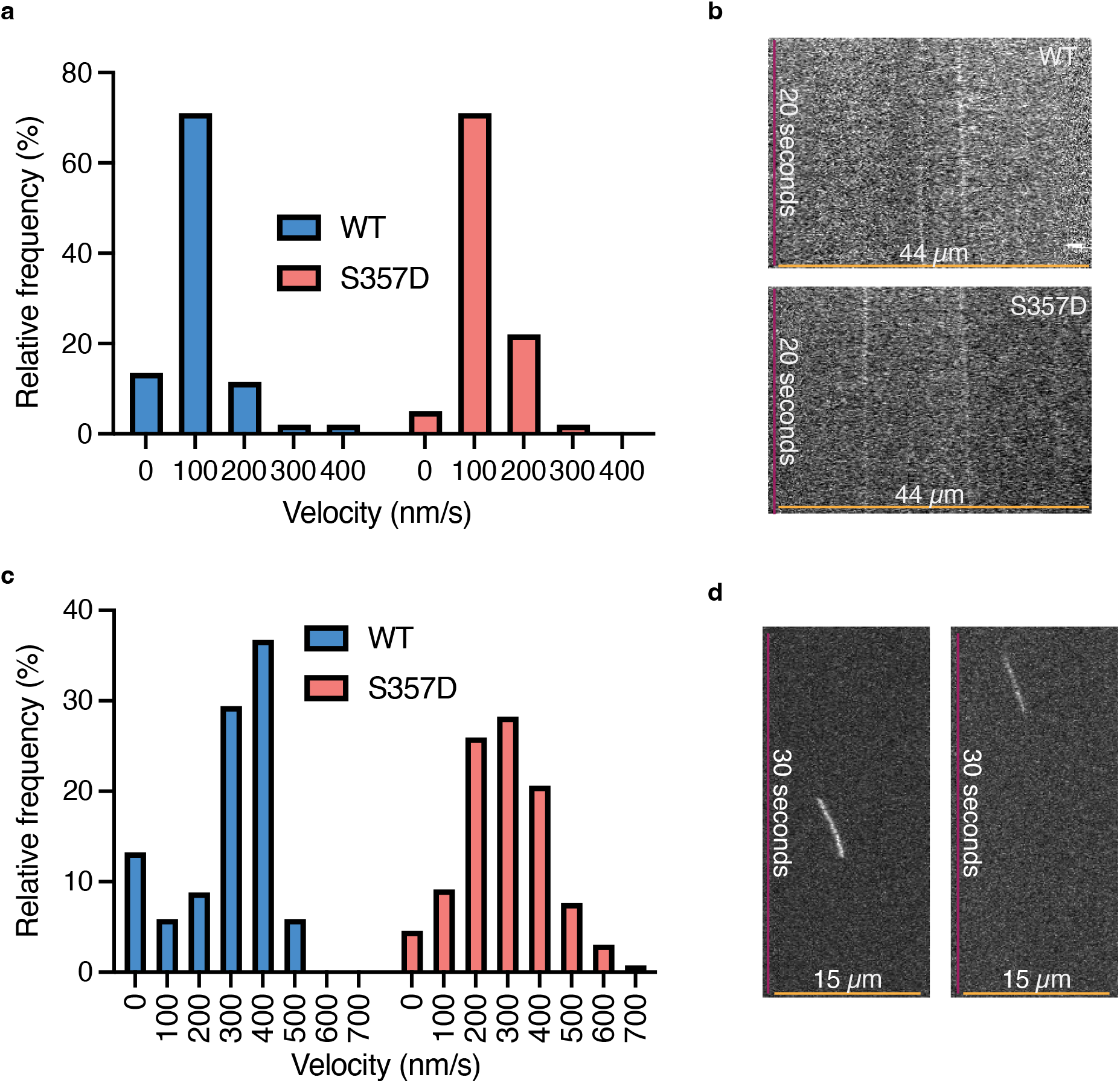
KIF18A S357D does not alter motor velocity. **(a)** Quantification of the relative frequency of velocity of KIF18A wild-type (WT) and S357D on taxol stabilized microtubules. Data were analyzed using a Kruskal Wallis with Dunn’s Test for Multiple Comparisons, and no statistical difference between WT and S357D was found. **(b)** Example kymographs depicting KIF18A wild-type (WT) and S357D motor movement on taxol stabilized microtubules. **(c)** Quantification of the relative frequency of velocity of KIF18A wild-type (WT) and S357D on dynamic microtubules. Data were analyzed using a Kruskal Wallis with Dunn’s Test for Multiple Comparisons, and no statistical difference between WT and S357D was found. **(d)** Example kymographs depicting KIF18A wild-type (WT) and S357D motor movement on dynamic microtubules.

### KIF18A S357D altered localization closely mimics localization patterns of a shortened neck-linker mutant (sNL)

KIF18A has an extended neck-linker (by 3 amino acids) compared to some other kinesins, including kinesin-1. This 17-residue structure is critical for promoting accumulation of KIF18A to the plus-ends of kinetochore microtubules by promoting navigation around microtubule associated proteins, which can serve as obstacles on the microtubule lattice (Malaby et al., 2019a). To determine if the KIF18A S357D mutation disrupts the ability of KIF18A’s neck-linker to promote plus-end accumulation via obstacle navigation, we compared the localization patterns of a shortened neck-linker (sNL) KIF18A mutant (Figure 8a) to that of KIF18A S357D in the presence and absence of the microtubule bundling protein HURP. We found that both KIF18A sNL and KIF18A S357D exhibited altered localization to peripheral microtubules and reduced accumulation at kinetochore microtubules in the presence of the microtubule bundling protein HURP (Figure 8b-c). Following HURP depletion, both the sNL and S357D KIF18A mutants displayed a modest increase in kinetochore microtubule localization (Figure 8b-c). However, both sNL and S357D mutants are unable to achieve kinetochore microtubule accumulation at levels comparable to wild-type KIF18A (Figure 8b-c). Given the similar localization patterns of the KIF18A sNL and S357D mutations, we hypothesize that KIF18A S357D may result in a shortened neck-linker like state.

**Figure 8:**
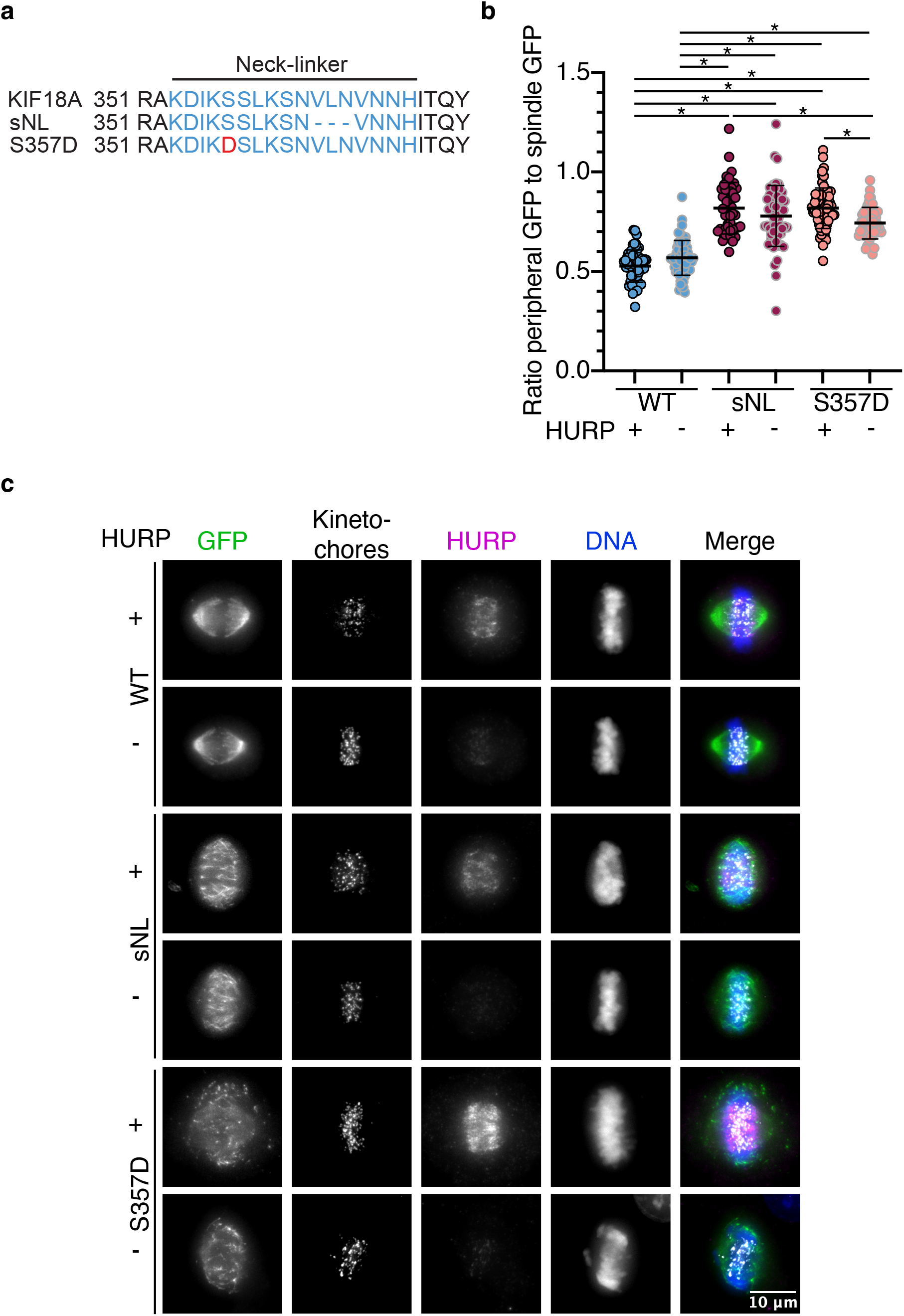
KIF18A S357D localization mimics that of a shortened neck-linker mutant. **(a)** KIF18A wild-type, shortened neck-linker (sNL), and S357D sequences. Blue text indicates neck-linker region. Red text indicates location of S357D mutation. **(b)** Quantification of GFP-KIF18A localization in periphery versus on kinetochore microtubules in mitotic cells. Solid horizontal line indicates mean, vertical lines indicate standard deviation. Each solid small dot represents a single cell. Black outlined dots indicate presence of HURP, gray outlined dots indicate HURP knockdown. Data were acquired from three experimental replicates and were analyzed using a one-way ANOVA with Tukey’s test for multiple comparisons. P value style: <0.05 (*), if no significance is indicated comparison was not significant. WT: wild-type. **(c)** Representative images of KIF18A localization in RPE1 cells. Cells were fixed approximately 24 hours after siRNA treatment to knockdown endogenous KIF18A and HURP and induction of GFP-KIF18A with doxycycline. Color indicates pseudo-color in merged image. +/- indicates presence or absence of HURP respectively. Brightness/contrast levels for HURP are set to be equivalent, brightness/contrast levels for kinetochores, DNA, and GFP are set differently to optimize visualization. Scale bar is 10 μm. WT: wild-type, sNL: shortened neck-linker, KD: knockdown.

## DISCUSSION

This body of work highlights how a single modification within the neck-linker of a kinesin can alter its sub-cellular localization and function. We have demonstrated that KIF18A S357D results in altered accumulation of the motor to peripheral microtubules of the spindle. The altered accumulation is also observed in monopolar mitotic cells and interphase cells, where an increase in KIF18A at the peripheral edge of a cell was observed. Surprisingly, despite this relocalization, the KIF18A S357D mutation only modestly reduces chromosome alignment and spindle length control. This was somewhat surprising since accumulation of KIF18A at the plus-ends of kinetochore microtubules is essential for its role in promoting chromosome alignment and maintaining spindle length (Stumpff et al., 2008; Malaby et al., 2019a; Du et al., 2010; Mayr et al., 2007). KIF18A promotes chromosome alignment by suppressing kinetochore microtubule plus-end dynamics (Stumpff et al., 2008, 2012; Du et al., 2010). Thus, we hypothesize that the small amount of KIF18A S357D that does accumulate at kinetochore microtubule plus-ends is sufficient to promote chromosome alignment and maintain spindle length, indicating that the level of KIF18A needed to promote chromosome alignment and spindle length maintenance may be relatively low.

In contrast, KIF18A S357D was unable to promote mitotic progression. KIF18A contributes to mitotic progression by promoting kinetochore-microtubule attachment and tension across kinetochores, which are necessary to satisfy the spindle assembly checkpoint (Janssen et al., 2018; Stumpff et al., 2012; Marquis et al., 2021; Zhu et al., 2005; Mayr et al., 2007; Czechanski et al., 2015). The fact that KIF18A S357D was unable to promote mitotic progression suggests that there is not sufficient KIF18A present at the plus-ends of kinetochore microtubules to allow for silencing of the spindle assembly checkpoint. Given the differential consequences of KIF18A S357D on promoting chromosome alignment and mitotic progression, future work is needed to discern the level of KIF18A necessary for each function. This question may be particularly relevant to understanding why chromosomally unstable tumor cells are sensitive to the loss of KIF18A, which is correlated with mitotic arrest (Marquis et al., 2021; Cohen-Sharir et al., 2021).

The most striking effect of expressing the KIF18A S357D mutant was in the observed mitotic spindle stabilization defects, which resulted in rotation of the mitotic spindle. Peripheral spindle microtubules play a critical role in contacting the cortex and maintaining mitotic spindle position (Garzon-Coral et al., 2016; Grill and Hyman, 2005). The most plausible explanation for this phenotype is that the accumulation of KIF18A S357D onto peripheral microtubules disrupts the interaction between the microtubules and the cell cortex, causing defects in maintaining spindle position and subsequent spindle rotation. However, we can’t rule out other possible scenarios where KIF18A S357D disrupts actin and myosin-dependent mechanisms that are necessary for maintaining spindle orientation (Fink et al., 2011).

Interestingly, the altered localization pattern of KIF18A S357D closely mimics that of a shortened neck-linker mutant, which may indicate that KIF18A S357D causes a kink in the neck-linker of KIF18A and effectively shortens this structure. In support of this idea, Alpha Fold predicts possible charge interactions between S357 and H10 (Jumper et al., 2021). Charge changes at S357 could, therefore, facilitate higher affinity interactions between S357 and H10 and lead to a shortened neck-linker-like state. Furthermore, expression of a shortened neck-linker mutant in HeLa cells led to similar effects on chromosome alignment as seen with the KIF18A S357D mutant, where sNL mutants were unable to promote chromosome alignment to the same extent as wild-type KIF18A but did not result in as severe of an unalignment phenotype as KIF18A knockdown alone (Malaby et al., 2019a).

The extended neck-linker of KIF18A allows for obstacle navigation and subsequent plus-end accumulation (Malaby et al., 2019a). When the neck-linker of KIF18A was shortened by three amino acids, the motor lost its canonical plus-end localization in HeLa cells, which was rescued by removal of the kinetochore microtubule bundling protein HURP. These data suggested that the altered localization of the sNL was caused by an inability of KIF18A to side-step around microtubule associated proteins in the absence of an extended neck-linker (Malaby et al., 2019a). One possible model for the altered accumulation of KIF18A S357D is that charge change at this site disrupts the ability of the motor to perform this necessary obstacle navigation, thereby excluding KIF18A S357D from the plus-end of the kinetochore microtubules and causing altered localization to peripheral microtubules. Kinetochore microtubules are known to recruit additional microtubule associated proteins (MAPs) compared to peripheral microtubules, including the microtubule bundling protein HURP (Silljé et al., 2006; Koffa et al., 2006). The additional MAPs on kinetochore microtubules may present an environment that sNL and S357D KIF18A are unable to navigate, resulting in accumulation on peripheral microtubules. While this idea is supported by the increased accumulation of KIF18A S357D and sNL at kinetochore microtubule plus-ends following HURP knockdown in RPE1 cells, this extent of accumulation did not match wild-type. This could be due to additional MAPs that serve as obstacles along kinetochore microtubules. HURP knockdown also had a stronger effect on sNL localization in HeLa cells than RPE1 cells, suggesting that the effects of individual MAPs on KIF18A navigation can vary by cell type (Malaby et al., 2019a).

In addition to causing changes in obstacle navigation, it is possible that the S357D mutation may alter KIF18A’s binding preference for the microtubule lattice. Subsets of microtubules within cells can differ in several ways. For example, the microtubule lattice can be either extended or compressed, which impacts the microtubule associated proteins that preferentially bind (Brouhard and Rice, 2018; Siahaan et al., 2022). Since neck-linker length affects the flexibility and coordination of the two motor heads in dimeric kinesins (Dogan et al., 2015; Shastry and Hancock, 2010; Hoeprich et al., 2014; Malaby et al., 2019a), it is possible that a shortened neck- linker or S357D mutation could alter the preference of KIF18A for compressed or extended lattices. If lattice compaction differs between kinetochore and peripheral microtubules within the spindle, perhaps this could contribute to KIF18A’s preference for peripheral microtubules. Given the changes in microtubule associated proteins that preferentially bind extended versus compressed lattices, it is also plausible that KIF18A S357D preferentially binds lattices which contain particular microtubule associated proteins or have fewer obstacles to navigate. In addition, post-translational modifications of microtubules impact the association and function of kinesins (Sirajuddin et al., 2014; Reed et al., 2006; Kaul et al., 2014; Barisic et al., 2015). Microtubule PTMs have the potential to impact charge interactions, and it is possible that the charge associated with KIF18A S357D alters the interaction of the motor with microtubule PTMs resulting in altered localization. While the sNL mutant does not have a net charge change, it is possible that charged residues that are normally available for interaction with the microtubule lattice are no longer accessible when the neck-linker is shortened, preventing typical charge interactions between KIF18A and the microtubule lattice. A similar mechanism may also be occurring with the KIF18A S357D mutant, whereby charge interactions between S357 and other regions of the motor hinder access to additional residues that would normally interact with the microtubule lattice. Given the similarities between the KIF18A S357D mutant and the shortened neck-linker mutant, it is possible that shortening the neck-linker of KIF18A directly or through the S357 mutation may alter the way the motor moves on microtubules containing PTMs.

Overall, this work highlights how a single PTM within the neck-linker of KIF18A causes altered accumulation to a different microtubule subpopulation in both mitotic and interphase cells. Similar to work focusing on modifications within the motor domain of kinesins (Muretta et al., 2018; Bickel et al., 2017; DeBerg et al., 2013) this altered accumulation upon modification may indicate a possible mechanism of regulation whereby PTMs in the neck-linker of kinesins contribute to their localization in a temporal and spatial dependent manner. One possible benefit for phosphorylation of KIF18A S357D in interphase may be to promote a subset of motor accumulating at the peripheral edge of a cell, where it could facilitate cellular functions including cell migration. It is also plausible that a subset of KIF18A is phosphorylated in mitosis, permitting differential regulation of the peripheral versus spindle microtubules, allowing for maintenance of spindle centering while also promoting chromosome alignment and mitotic progression. Future work will focus on understanding the precise mechanism that promotes the altered accumulation of KIF18A 357D to peripheral spindle microtubules.

## MATERIALS AND METHODS

### Cell Culture and Transfections

HeLa Kyoto and RPE1 acceptor cells were a kind gift from Ryoma Ohi, University of Michigan. Cells were cultured at 37°C with 5% CO2 in MEM-α medium (Gibco, 12561072) containing 10% Fetal Bovine Serum (FBS) (Gibco, 16000044). HeLa Kyoto and RPE1 acceptor cells for recombination were maintained in Blasticidin (Thermo Fisher Scientific, R21001). Once inducible clones were established by recombination, HeLa Kyoto inducible cell lines were maintained in 1 μg/mL puromycin (Thermo Fisher Scientific, A11138-03) and RPE1 inducible lines were maintained in 5 μg/mL puromycin (Thermo Fisher Scientific, A11138-03).

For siRNA transfections conducted in a 24-well plate 5pmol siRNA was incubated in Opti-MEM Reduced Serum Media (Gibco, 31985062) with Lipofectamine RNAiMax Transfection Reagent (Invitrogen, 13778150) for 5 minutes at room temperature before adding dropwise to wells. KIF18A siRNA used was a 1:1 mixture of the following two Silencer Select Validated siRNAs (5’ to 3’ sequence): GCUGGAUUUCAUAAAGUGGtt (Ambion, AM51334) CGUUAACUGCAGACGUAAAtt (Ambion, 4392420). For control knockdown a 1:1 mixture of Silencer Select Negative Control #2 (Ambion, 4390846) and Silencer Select Negative Control #1 (4390843) was used. For experiments with HURP knockdown the custom siRNA with 5’ to 3’ sequence used was AGUUACACCUGGACUCCUUTT (Qiagen, 1027423).

### Immunofluorescence

For fixed cell immunofluorescence experiments cells were seeded on acid-treated 12mm glass coverslips in a 24-well dish and fixed in -20oC methanol (Fisher Scientific, A412-1) for 3 minutes, -20oC methanol with 1% paraformaldehyde (Electron Microscopy Sciences, 15710) for 10 minutes, or 1X BRB80, 4mM EGTA (Sigma Aldrich, E4378) 0.5% Triton-X (Sigma Aldrich, 93443) (MTSB) with 0.5% glutaraldehyde (Electron Microscopy Sciences, 16360) for 10 minutes. Glutaraldehyde fixed cells were subsequently quenched with 0.1% sodium borohydride (Fisher Scientific, S67825) in 1X Tris-Buffered Saline (TBS; 150 mM NaCl, 50 mM Tris base, pH 7.4) for 15 minutes two times. After fixation coverslips were washed 2-3 times in 1X TBS for minutes each. Coverslips were then blocked with 20% goat serum in antibody dilution buffer (Abdil: TBS pH 7.4, 1% Bovine Serum Albumin (BSA), 0.1% Triton X-100, and 0.1% sodium azide) for 1 hour at room temperature. Prior to incubation with primary antibodies, coverslips were washed two times in 1X TBS for 5 minutes. Primary antibodies were diluted 1:1 in glycerol and stored at -20oC. Prior to incubation on coverslips primary antibodies were further diluted in Abdil at indicated concentrations and incubated for the indicated duration. Prior to secondary antibody incubation coverslips were washed two additional times for 5 minutes in 1X TBS. The following secondary antibodies conjugated to AlexaFluor 488, 594, and 647 were diluted 1:1 in glycerol and stored at -20oC prior to diluting 1:500 in abdil for incubation on coverslips for 1 hour at room temperature (Invitrogen Molecular Probes A11007, A11013, A11014, A11029, A11032, A11034, A11037, A11076, A21236, A21245, A21247, A21450, A32931,). After secondary antibody incubation coverslips were washed an additional two times in 1X TBS for 5 minutes at room temperature prior to mounting in Prolong Gold anti-fade mounting medium with DAPI (Invitrogen Molecular Probes, P36935).

### Microscopy

All fixed and live cell images or movies were acquired on a Ti-E inverted microscope (Nikon Instruments) or on a Ti-2E inverted microscope (Nikon Instruments) both driven by NIS Elements (Nikon Instruments). Images were captured using either a Clara cooled charge-coupled device (CCD) camera (Andor) or Prime BSI scientific complementary metal-oxide-semiconductor (sCMOS) camera (Teledyne Photometrics) with a Spectra-X light engine (Lumencore). Imaging was conducted with the following Nikon objectives: Plan Apo 40X 0.95 numerical aperture (NA), Plan Apo λ 60X 1.42 NA, and APO 100X 1.49 NA. Both microscopes contain environmental chambers held at 37°C. For live cell-imaging, cells were imaged in CO2-independent media (Gibco #18045-088) supplemented with 10% fetal bovine serum (Gibco #16000-044). All images were processed and analyzed using ImageJ/Fiji (Schneider et al., 2012; Schindelin et al., 2012).

TIRF microscopy experiments with taxol stabilized microtubules were conducted at 37oC using a Ti-E inverted microscope (Nikon Instruments) equipped with an EMCCD camera (Andor), 488 nm and 567 nm lasers, and a 100X Apo TIRF (1.49 numerical aperture) objective lens (Nikon). TIRF microscopy experiments with dynamic microtubules were conducted at 37oC using an inverted Eclipse Ti-E microscope (Nikon) with dual iXon Ultra Electron Multiplying charge-coupled device cameras and a 100XApo TIRF objective lens (1.49 numerical aperture) driven by NIS Elements, version 4.51.0.

### Generation and validation of HeLa Kyoto and RPE1 inducible cell lines

Both HeLa Kyoto and RPE1 inducible cell lines were generated using methods described previously (Khandelia et al., 2011). In brief, a wild-type KIF18A siRNA and puromycin resistant plasmid containing LoxP sites was generated for recombination mediated cassette exchange. The resulting plasmid was transfected into either HeLa Kyoto or RPE1 acceptor cells (Sturgill et al., 2016) with recombinase using an LTX transfection (Thermo Fisher Scientific). HeLa Kyoto cells that had undergone exchange were selected for with 1 μg/mL puromycin for 48 hours followed by a stricter selection with 2 μg/mL puromycin for 48 hours prior to switching back to 1 μg/mL puromycin. Prior to transfection for recombination RPE1 acceptor cells were seeded as single cells, clones were grown up, and then selected for based on susceptibility to puromycin treatment. The clones which were most susceptible were used for recombination and selected for with 10 μg/mL puromycin for 72 hours followed by a stricter selection with 20 μg/mL puromycin for 72 hours prior to switching back to 10 μg/mL. The resulting KIF18A inducible cell lines were maintained in MEM Alpha (Life Technologies) with 10% FBS (Life Technologies) and 1 μg/mL puromycin (HeLa Kyoto) or 5 μg/mL puromycin (RPE1) at 37°C, 5% CO2.

KIF18A wild-type (Czechanski et al., 2015), S357A (generated from a synthesized fragment from ThermoFisher Scientific and mutagenesis), and S357D (generated through mutagenesis) siRNA resistant fragments, and pEM791 vector (Sturgill et al., 2016) were amplified with primers designed for Gibson Assembly (New England BioLabs) and subsequently assembled via Gibson Assembly. All plasmids were sequenced to confirm sequence identify prior to recombination.

Genomic DNA was extracted from resulting HeLa Kyoto and RPE1 inducible cell lines (QIAmp DNA Blood Mini Kit Qiagen #51106) and sequenced through the S357 mutation site (Eurofins) to verify correct incorporation of the desired construct. Inducible constructs were expressed by the addition of 2 μg/mL doxycycline (Thermo Fisher Scientific #BP26531).

### KIF18A expression level quantification in HeLa Kyoto and RPE1 inducible cell lines

For quantification of KIF18A expression by immunofluorescence, cells were seeded on 12mm coverslips and endogenous KIF18A was depleted 24 hours later as described above. Simultaneously, expression of GFP-KIF18A constructs was induced with the addition of 2 μg/mL doxycycline. 24 hours post-knockdown and induction coverslips were fixed in -20oC methanol for 3 minutes and stained for ms-gamma tubulin 1:500 (Sigma Aldrich, T5326), guinea-pig CENP-C 1:25 (MBL, PD030), and rabbit KIF18A (Bethyl, A301-080A). All primary antibodies were incubated for 1 hour at room temperature with shaking. Random mitotic cells were imaged from each condition. Expression levels were quantified in ImageJ by drawing an ROI centered around but larger than the mitotic spindle and measuring KIF18A intensity. Background levels for each cell were measured by moving the ROI to an area with no cells and measuring the KIF18a intensity. The mean background signal for each image was subsequently subtracted from the mean spindle KIF18A signal. All values were then normalized to the mean background subtracted control KIF18A intensity. Mean and standard deviations are reported for three individual biological replicates. The total number of cells analyzed for each condition was RPE1 control = 61 cells, RPE1 KIF18A knockdown = 62 cells, RPE1 GFP = 64 cells, RPE1 WT = 61 cells, RPE1 S357A = 60 cells, and RPE1 S357D = 64 cells, HeLa Kyoto control = 57 cells, HeLa Kyoto KIF18A knockdown = 62 cells, HeLa Kyoto GFP = 65 cells, HeLa Kyoto WT = 61 cells, HeLa Kyoto S357A = 61 cells, and HeLa Kyoto S357D = 64 cells.

### KIF18A localization imaging

For images displaying KIF18A localization in mitosis, cells were seeded on 12mm coverslips. Endogenous KIF18A was depleted 24 hours later as described above, and exogenous GFP-KIF18A was induced simultaneously with the addition of 2 μg/mL doxycycline. 24 hours after induction of GFP-KIF18A and endogenous KIF18A knockdown cells were fixed for 3 minutes in - 20oC methanol and subsequently stained for rabbit GFP 1:500 (Invitrogen, A11122) for one hour at room temperature with shaking.

For visualization of KIF18A localization in monopolar spindles cells were seeded on 12mm coverslips. Endogenous KIF18A was depleted 24 hours later as described above, and exogenous GFP-KIF18A was induced simultaneously with the addition of 2 μg/mL doxycycline. 24 hours after induction and knock down cells were treated with 100μM monastrol for 2-3 hours to cause spindle collapse prior to fixing in -20oC methanol for 5 minutes. Cells were then subsequently stained for rabbit-gamma-tubulin 1:250 (Sigma Aldrich, T5192), mouse-EB1 1:50 (BD Transduction Laboratories, 610535), and chicken-GFP 1:500 (Invitrogen, A10262) all for one hour at room temperature with shaking.

For images displaying interphase localization cells were seeded in a glass bottom 35mm dish. Endogenous KIF18A was depleted 24 hours later as described above, and exogenous GFP- KIF18A was induced simultaneously with the addition of 2 μg/mL doxycycline. 24 hours after induction media was exchanged to CO2 Independent Media (Gibco, 18045088) with 50nM siR-tubulin (Cytoskeleton, cy-sc-002), 1:20,000 spy595 DNA (Cytoskeleton, cy-sc-301), 10uM verapamil (Cytoskeleton, cy-sc-002), and 2 μg/mL doxycycline and left to incubate at 37 oC for at least 2 hours prior to imaging.

### Quantification of chromosome alignment and spindle length

For chromosome alignment and spindle length analysis cells were seeded on 12mm coverslips. Endogenous KIF18A was depleted 24 hours later as described above, and exogenous GFP-KIF18A was induced simultaneously with the addition of 2 μg/mL doxycycline. 24 hours after induction of GFP-KIF18A and endogenous KIF18A knockdown cells were fixed for 3 minutes in - 20oC methanol and subsequently stained for ms-gamma-tubulin 1:500 (Sigma Aldrich, T5326), guinea-pig-CENP-C 1:250 (MBL, PD030) and rabbit-GFP 1:500 (Invitrogen, A11122). All primary antibodies were incubated on coverslips for one hour at room temperature with shaking. Approximately twenty random metaphase mitotic cells in which both poles of the mitotic spindle were in focus in the same z plane for each condition were imaged. As described previously, (Fonseca and Stumpff, 2016; Malaby et al., 2019b), a line with length defined by the distance between spindle poles was used to measure the distribution of CENP-C labeled kinetochores using the Plot Profile command in ImageJ/Fiji. The CENP-C signal intensity was normalized internally to its highest value and plotted as a function of distance along the pole-to-pole axis using a custom macro in MatLab. The plots were then fitted to a Gaussian curve and the Full-Width at Half Maximum (FWHM), and the spindle length are reported for each cell analyzed. Mean and standard deviations are reported from three biological replicates for each condition. A one-way ANOVA with Tukey’s Test for Multiple Comparisons was conducted for statistical analysis. For each HeLa Kyoto condition the following cell numbers were analyzed control = 60 cells, KIF18A knockdown = 56 cells, GFP = 61 cells, WT = 64 cells, S357A = 66 cells, and S357D = 67 cells. For each RPE1 condition the following cell numbers were analyzed control = 67 cells, KIF18A knockdown = 88 cells, GFP = 88 cells, WT = 66 cells, S357A = 69 cells, and S357D = 67 cells.

### Analysis of mitotic timing and arrest

To determine mitotic timing, cells were seeded in glass-bottom 24-well dishes. 24 hours later endogenous KIF18A was knocked down as described above and exogenous KIF18A was induced with 2 μg/mL doxycycline. 24 hours after induction and knock down media was exchanged to CO2 Independent Media (Gibco, 18045088) supplemented with 2 μg/mL doxycycline for induced conditions. Multiple fields of view for each condition were imaged overnight at 40X magnification, imaging every 3 minutes. To quantify mitotic timing the timepoint at nuclear envelope breakdown and anaphase onset was determined and the difference was calculated to determine time from nuclear envelope breakdown to anaphase onset. Cell that entered mitosis at least one hour prior to the end of mitotic timing movies but never entered anaphase were quantified as cells that never divide. For mitotic timing, median values are reported from three biological replicates for each condition. A Kruskal Wallis Test with Dunn’s Test for Multiple Comparisons was conducted for statistical analysis. For each HeLa Kyoto condition the following cell numbers were analyzed control = 231 cells, KIF18A knockdown = 136 cells, GFP = 154 cells, WT = 162 cells, S357A = 182 cells, and S357D = 177 cells. For each RPE1 condition the following cell numbers were analyzed control = 114 cells, KIF18A knockdown = 95 cells, GFP = 134 cells, WT = 94 cells, S357A = 99 cells, and S357D = 86 cells. For mitotic arrest the percent of cells that never divide is reported from three experimental replicates.

### Quantification of angle between spindle poles and spindle rotation

For analysis of the angle between spindle poles cells were seeded on 12mm coverslips. Endogenous KIF18A was depleted 24 hours later as described above, and exogenous GFP-KIF18A was induced simultaneously with the addition of 2 μg/mL doxycycline. 24 hours after induction of GFP-KIF18A and endogenous KIF18A knockdown cells were fixed for 3 minutes in -20oC methanol and subsequently stained for rabbit-KIF18A 1:100 (Bethyl, A301-080A), guinea-pig- CENP-C (MBL, PD030), and mouse-gamma-tubulin (Sigma Aldrich, T5326). All primary antibodies were incubated on coverslips for one hour at room temperature with shaking. Approximately 20 random metaphase mitotic cells per condition were imaged at 60X magnification with 41 z stacks recorded at 0.25 μm/step. Using the point tool in ImageJ the two poles of each cell were selected at the z plane where they were in focus and the x, y, and slice information was recorded. A custom macro in ImageJ was created to determine the angle between the two poles for each cell by using the dot product. The mean and standard deviations of the pole angle are reported for three biological replicates for each condition. A one-way ANOVA with Tukey’s Test for Multiple Comparisons was conducted for statistical analysis. For each RPE1 condition the following cell numbers were analyzed control = 81 cells, KIF18A knockdown = 80 cells, GFP = 86 cells, WT = 85 cells, S357A = 82 cells, and S357D = 86 cells.

For quantification of spindle rotation cells were seeded in glass-bottom 24-well dishes. 24 hours later endogenous KIF18A was knocked down as described above and exogenous KIF18A was induced with 2 μg/mL doxycycline. 24 hours after induction and knock down media was exchanged to CO2 Independent Media (Gibco, 18045088) supplemented with 2 μg/mL doxycycline for induced conditions. Multiple fields of view per condition were imaged every minute for 30 minutes with 21 z stacks taken at 0.25 μm/step. To quantify spindle rotation one spindle pole of a cell was tracked using MTrackJ in ImageJ and the subsequent track was measured. From track measurements the displacement in microns per minute was calculated. The mean and standard deviations from three biological replicates are reported for each condition. A one-way ANOVA with Tukey’s Test for Multiple Comparisons was conducted for statistical analysis. For each RPE1 condition the following cell numbers were analyzed WT = 12 cells, S357A = 16 cells, and S357D = 24 cells.

### Analysis of polymerizing microtubules and tubulin density

For quantification of EB1 to determine changes in microtubule polymerization cells were seeded on 12mm coverslips. Endogenous KIF18A was depleted 24 hours later as described above, and exogenous GFP-KIF18A was induced simultaneously with the addition of 2 μg/mL doxycycline. 24 hours after induction and knock down cells were treated with 100μM monastrol for 2-3 hours to cause spindle collapse prior to fixing in -20oC methanol for 5 minutes. Cells were then subsequently stained for rabbit-gamma-tubulin 1:250 (Sigma Aldrich, T5192), mouse-EB1 1:50 (BD Transduction Laboratories, 610535), and chicken-GFP 1:500 (Invitrogen, A10262) all for one hour at room temperature with shaking. Approximately 20 random mitotic cells were imaged per condition at 60X magnification with 13 z slices taken at 0.2 μm/step. The ratio of peripheral to spindle EB1 or GFP was determined by drawing an ROI that encompassed the DNA and measuring the GFP and EB1 intensity in that ROI. A 2.5-micron band was then created around the DNA ROI and the EB1 and GFP intensity within that band was measured. The ratio between the two ROIs was then calculated. The mean and standard deviation from three biological replicates is reported. A one-way ANOVA with Tukey’s Test for Multiple Comparisons was conducted for statistical analysis. For each condition the following cell numbers were analyzed WT = 46 cells, S357A = 49 cells, and S357D = 56 cells.

For analysis of microtubule density cells were seeded on 12mm coverslips. Endogenous KIF18A was depleted 24 hours later as described above, and exogenous GFP-KIF18A was induced simultaneously with the addition of 2 μg/mL doxycycline. 24 hours after induction and knock down cells were fixed in 0.5% glutaraldehyde in MTSB for 10 minutes and then quenched two times for 15 minutes each with 0.1% sodium borohydride (Fisher Scientific, S67825) in 1X TBS. Cells were then subsequently stained for human-anti-centromere (ACA) 1:200 (Antibodies Inc., 15-235), ms-DM1-alpha 1:150 (Sigma Aldrich, T6199), and rabbit-GFP 1:400 (Invitrogen, A11122) all overnight at 4oC with shaking. Approximately 20 random metaphase mitotic cells were imaged per condition at 60X magnification with 41 z slices taken at 0.2 μm/step. The mean spindle tubulin intensity was determined by drawing an ROI that encompassed the kinetochore microtubules of the mitotic spindle and measuring the tubulin intensity in that ROI. A 4.5-micron band was then created around the spindle ROI and the tubulin intensity within that band was measured. The background tubulin intensity was measured with an ROI in a region containing no cells and the subsequent measurement was subtracted from both the mean spindle and mean peripheral tubulin intensity for a given cell. The median from a minimum of three biological replicates is reported. A Kruskal Wallis Test with Dunn’s Test for Multiple Comparisons was conducted for statistical analysis. For each condition the following cell numbers were analyzed control = 85 cells, KIF18A knockdown = 84 cells, GFP = 84 cells, WT = 80 cells, S357A = 63 cells, and S357D = 83 cells.

### smTIRF to quantify KIF18A velocity

For analysis of single molecule velocity GFP-KIF18A was isolated as cell lysates by seeding in a 100mm dish and inducing expression of exogenous GFP-KIF18A with the addition of 2 μg/mL doxycycline. Cells were arrested in mitosis with the addition of 100 ng/mL nocodazole overnight prior to obtaining cell pellet. The cell pellet was resuspended in lysis buffer (300 mM KCl, 1 mM MgCl2, 0.1 mM ATP, 1X Roche Protease/PPase inhibitor, and 10% glycerol in 1X BRB80) for 15 minutes on ice prior to shearing DNA by passing through a narrow-gauge (No 27) hypodermic needle. The sheared solution was then left on ice for an additional 15 minutes before spinning at 16,000 RCF in a microcentrifuge at 4°C for 20 minutes. The cleared lysate (supernatant) was snap frozen in liquid nitrogen and stored at -80°C prior to use.

For analysis on taxol stabilized microtubules, rhodamine labeled microtubules (Cytoskeleton, bovine) stabilized with 20 μM taxol (Selleck Chemicals, S1150) were prepared with a 100:1 ratio of unlabeled GTP tubulin to rhodamine labeled tubulin. The mixture of unlabeled and labeled tubulin was allowed to polymerize at 37oC for 20 minutes prior to stabilizing with 20 μM paclitaxel in DMSO (Selleck Chemicals, S1150). Experiments were conducted in silanized flow cells. Silanized slides (Deckgläser Microscope Coverglass, 24 x 60 mm, 170 ± 5 µm, No. 1.5H) were prepared by first washing in 100% methanol for 2 hours shaking at room temperatures. Slides were subsequently dried with nitrogen and cleaned with a UV-ozone plasma (Harrick plasma cleaner) for 3 minutes. Slides were then incubated in 99.2% toluene (Sigma-Aldrich, 244511), 0.173% 2-methoxy(polyethyleneoxy)propyltrimethoxysilane (Gelest, SIM6492.7) and 0.62% n-butylamine (Acros Organics) with flowing nitrogen gas for 90 minutes. Coverslips were subsequently washed twice in toluene and dried with nitrogen gas before curing for 30 minutes with flowing nitrogen gas. Flow cells chambers were assembled using silanized coverslips adhered to a 25 X 75mm glass slide (Thermo Scientific, 3011-002) with two pieces of double-sided tape which resulted in a ∼20μl chamber. Microtubules were linked to PEG-silanized coverslips by incubating chambers with 33 μg/mL monoclonal anti-beta-tubulin III (neuronal) antibody (Sigma-Aldrich, T8578) for 5 minutes prior to washing with 10 mg/mL BSA for 2 minutes. Flow chambers were then incubated with 1 μM assembled microtubules for 10 minutes. Experiments were conducted in 1X BRB80 containing 10mM DTT, 100mM KCl, and 1X Oxygen Scavenger Mix (OSM; 0.045 mg/mL catalase, 0.066 mg/mL glucose oxidase, and 5.8 mg/mL glucose) in 20μM paclitaxel. Images were collected at 10 frames/sec. Motor velocity was calculated by creating kymographs from individual microtubules in ImageJ, creating a rectangle that bound the sloped line, and measuring the width (run length in pixels) and height (number of time frames) of the resulting rectangle. The subsequent velocity was calculated by multiplying the width (in pixels) by the pixel length and dividing this by the time of the run (height multiplied by the time interval in seconds). For each condition the number of motor runs analyzed was WT = 52, S357D = 106.

For analysis on dynamic microtubules rhodamine labeled microtubules (bovine) were prepared as described previously (Cario et al., 2022b). Experiments were conducted in flow cells and microtubules were linked to PEG-silanized coverslips as described previously (Cario et al., 2022a; Stern et al., 2017). Experiments were conducted in 1X BRB80 containing 10mM DTT, 100mM KCl, and 1X Oxygen Scavenger Mix (OSM; 0.045 mg/mL catalase, 0.066 mg/mL glucose oxidase, and 5.8 mg/mL glucose). Images were collected at 10 frames/sec. Motor velocity was calculated by creating kymographs from individual microtubules in ImageJ, creating a rectangle that bound the sloped line, and measuring the width (run length in pixels) and height (number of time frames) of the resulting rectangle. The subsequent velocity was calculated by multiplying the width (in pixels) by the pixel length and dividing this by the time of the run (height multiplied by the time interval in seconds). For each condition the number of motor runs analyzed was WT = 68, S357D = 131.

### Analysis of KIF18A S357 mutants and KIF18A sNL mutant localization in the presence and absence of HURP

To analyze GFP-KIF18A localization in the presence and absence of HURP cells were seeded on 12mm coverslips. Endogenous KIF18A and HURP was depleted 24 hours later as described above, and exogenous GFP-KIF18A was induced simultaneously with the addition of 2 μg/mL doxycycline. 24 hours after induction cells were treated with 25 μM MG132 for 1-2 hours prior to fixing in 1% paraformaldehyde in -20oC methanol for 10 minutes. Coverslips were subsequently stained for chicken-GFP 1:500 (Invitrogen, A10262), guinea-pig-CENP-C 1:250 (MBL, PD030), and rabbit-HURP 1:500 (Bethyl, A300-851A), all for one hour at room temperature with shaking. Approximately 20 random metaphase mitotic cells were imaged for each condition per biological replicate at 60X magnification with 31 z stacks taken at 0.25 μm/step. To quantify the ratio of peripheral GFP-KIF18A to spindle GFP-KIF18A z projected images were rotated so the two spindle poles were horizontal. A 20-pixel wide line twice the length of the distance of the kinetochore mass was centered over the kinetochores and the distribution of GFP-KIF18A was measured using the Plot Profile command in ImageJ. The resulting intensity data was normalized so the line went from zero to one. The average intensity corresponding with the top and bottom quarters of the line (0-0.25 and 0.74-1) was determined and recorded as the peripheral GFP- KIF18A intensity. The average intensity of GFP-KIF18A corresponding to the middle half of the line (0.26-0.75) was determined and recorded as the average spindle GFP-KIF18A intensity. The ratio between peripheral and spindle GFP-KIF18A was then calculated by dividing the peripheral average by the spindle average. A one-way ANOVA with Tukey’s Test for Multiple Comparisons was conducted for statistical analysis. For each condition the following cell numbers were analyzed: WT with HURP = 59 cells, WT HURP KD = 64 cells, sNL with HURP = 46 cells, sNL HURP KD = 55 cells, S357D with HURP = 66 cells, S357D HURP KD = 50 cells.

## Acknowledgments

This work was supported by NIH R01 GM121491 to JS, NIH R01GM130556 to JS, NIH R35 GM144133 to JS, NIH R01 GM132646 to CB, and NSF GRF 1842491 to KAQ. We thank Ryoma Ohi for acceptor cells and reagents used for recombination mediated cassette exchange. We thank Alex Thompson and Sarah Vandal for generation of the HeLa Kyoto and RPE1 GFP cell lines and for assistance in generating the HeLa Kyoto wild-type KIF18A cell line.

## Video Legends

**Supplementary Video 1: Stabilization of the microtubules of the mitotic spindle with taxol results in additional accumulation of wild-type KIF18A at kinetochore microtubule plus-ends.** Live-cell imaging of RPE1 wild-type GFP-KIF18A inducible cell line. Cells were imaged approximately 24 hours after siRNA treatment to knockdown endogenous KIF18A and induction of GFP-KIF18A with doxycycline. Immediately prior to imaging, cells were spiked with 10 μM paclitaxel.

**Supplementary Video 2: Stabilization of the microtubules of the mitotic spindle with taxol results in additional accumulation of S357A KIF18A at kinetochore microtubule plus-ends.** Live-cell imaging of RPE1 S357A GFP-KIF18A inducible cell line. Cells were imaged approximately 24 hours after siRNA treatment to knockdown endogenous KIF18A and induction of GFP-KIF18A with doxycycline. Immediately prior to imaging, cells were spiked with 10 μM paclitaxel.

**Supplementary Video 3: Stabilization of the microtubules of the mitotic spindle with taxol does not result in additional accumulation of S357D KIF18A at kinetochore microtubule plus-ends.** Live-cell imaging of RPE1 S357D GFP-KIF18A inducible cell line. Cells were imaged approximately 24 hours after siRNA treatment to knockdown endogenous KIF18A and induction of GFP-KIF18A with doxycycline. Immediately prior to imaging, cells were spiked with 10 μM paclitaxel.

**Supplementary Video 4: Wild-type KIF18A maintains spindle centering during mitosis.** Live-cell imaging of GFP-KIF18A wild-type in mitosis. Cells were imaged approximately 24 hours after siRNA treatment and induction of GFP-KIF18A with doxycycline and 3 hours after addition of siR-tubulin to label microtubules.

**Supplementary Video 5: S357D KIF18A displays defects in spindle centering in mitosis.** Live-cell imaging of GFP-KIF18A S357D in mitosis. Cells were imaged approximately 24 hours after siRNA treatment and induction of GFP-KIF18A with doxycycline and 3 hours after addition of siR-tubulin to label microtubules.

**Supplementary Video 6: In the absence of siR-tubulin wild-type KIF18A still maintains spindle positioning.** Live-cell imaging of GFP-KIF18A wild-type in mitosis. Cells were imaged approximately 24 hours after siRNA treatment and induction of GFP-KIF18A with doxycycline.

**Supplementary Video 7: In the absence of siR-tubulin S357D KIF18A is still unable to maintain spindle positioning.** Live-cell imaging of GFP-KIF18A S357D in mitosis. Cells were imaged approximately 24 hours after siRNA treatment and induction of GFP-KIF18A with doxycycline.

---

**Supplementary Figure 1:**
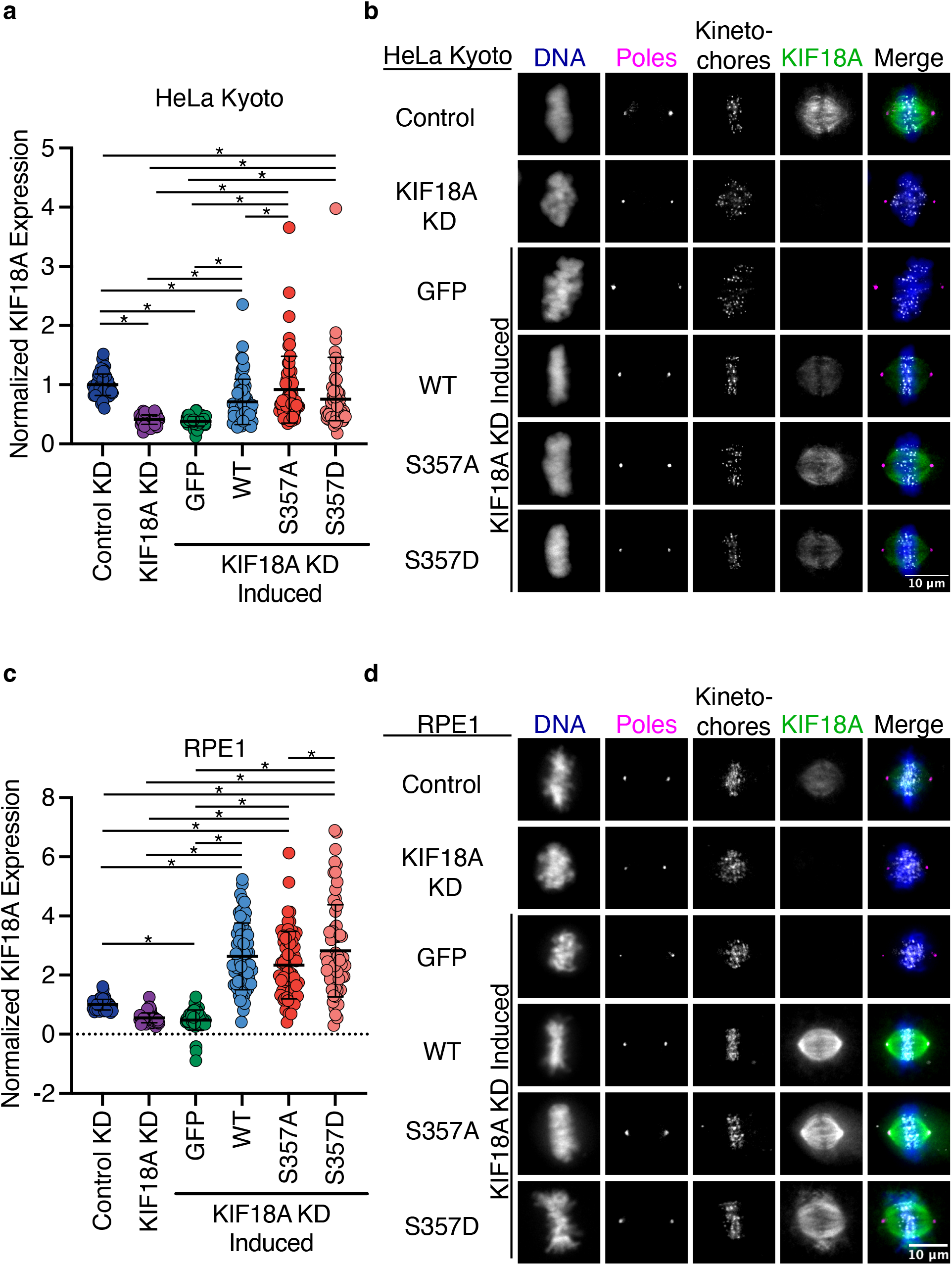
KIF18A inducible cell lines have similar expression. **(a)** Quantification of KIF18A expression in HeLa Kyoto cells from KIF18A antibody intensity. Fluorescence values were normalized to the mean control KIF18A intensity. Solid horizontal line indicates mean, vertical lines indicate standard deviation. Each dot represents a single cell. Data were acquired from three experimental replicates. Data were analyzed using a one-way ANOVA with Tukey’s test for multiple comparisons. P value style: <0.05 (*), if no significance is indicated result was not significant ( > 0.05). **(b)** Representative immunofluorescence images of KIF18A expression in HeLa Kyoto cells. Cells were fixed approximately 24 hours after siRNA treatment to knockdown endogenous KIF18A and induction of GFP-KIF18A with doxycycline. Colors indicate pseudo-color in merged image. Brightness/contrast levels for KIF18A are set to be equivalent across conditions. Brightness/contrast levels for DNA, poles, and kinetochores are set differently to optimize visualization. Scale bar is 10 μm. WT: wild-type. KD: knockdown. **(c)** Quantification of KIF18A expression in RPE1 cells from KIF18A antibody intensity. Fluorescence values were normalized to the mean control KIF18A intensity. Solid horizontal line indicates mean, vertical lines indicate standard deviation. Each dot represents a single cell. Data were acquired from three experimental replicates. Data were analyzed with a one-way ANOVA with Tukey’s test for multiple comparisons. P value style: <0.05 (*), if no significance is indicated result was not significant ( > 0.05). **(d)** Representative immunofluorescence images of KIF18A expression in RPE1 cells. Cells were fixed approximately 24 hours after siRNA treatment to knockdown endogenous KIF18A and induction of GFP-KIF18A with doxycycline. Colors indicate pseudo-color in merged image. Scale bar is 10 μm. WT: wild-type. KD: knockdown.

**Supplementary Figure 2:**
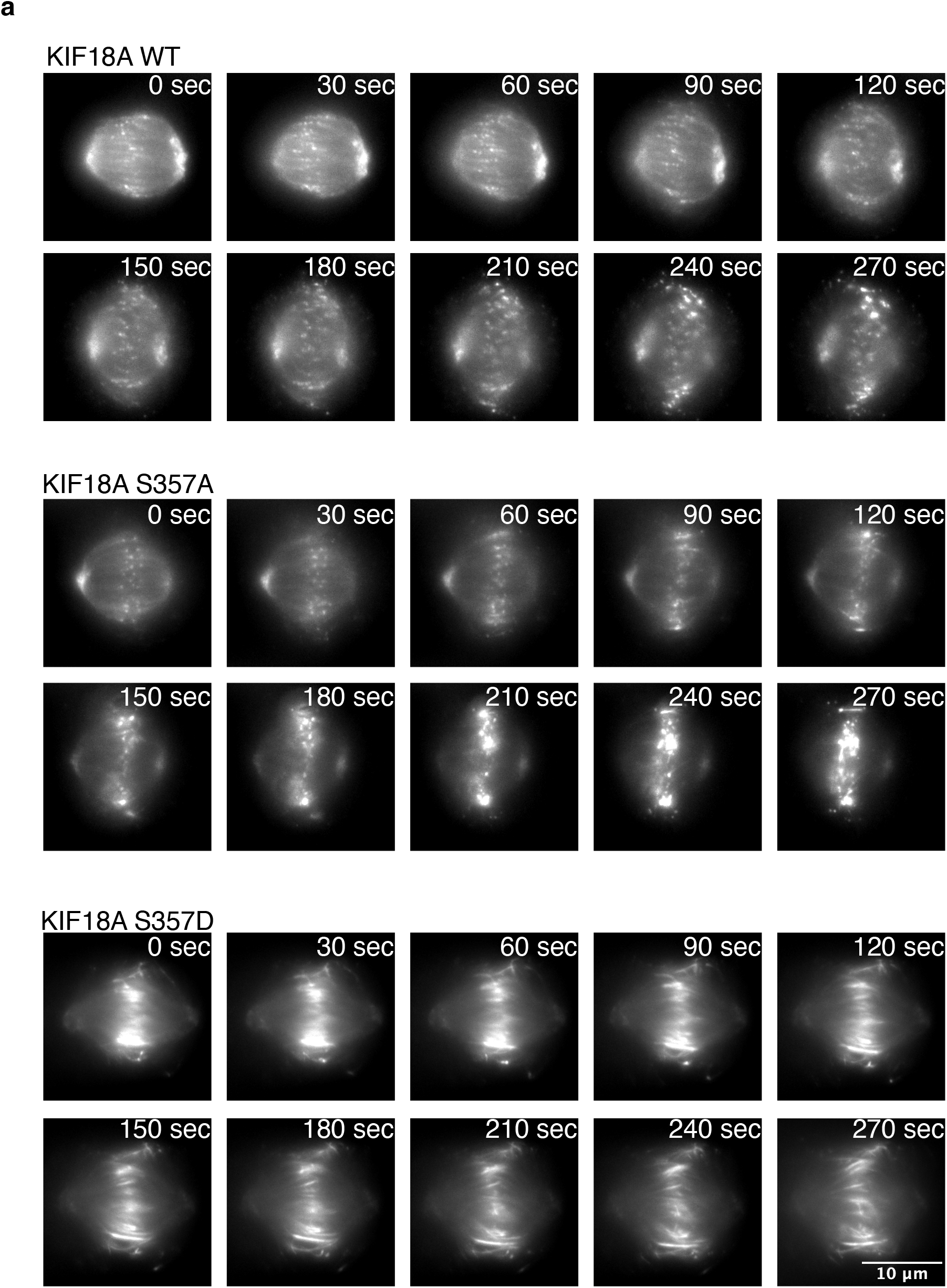
Stabilization of microtubules with taxol does not rescue peripheral localization of KIF18A S357D. **(a)** Representative still images from live-cell imaging of RPE1 GFP- KIF18A inducible cells lines. Cells were imaged approximately 24 hours after siRNA treatment to knockdown endogenous KIF18A and induction of GFP-KIF18A with doxycycline. Immediately prior to imaging, cells were spiked with 10 μM paclitaxel (0 sec) and time stamps indicate elapsed time after taxol treatment. Scale bar is 10 μm. WT: wild-type.

**Figure S3:**
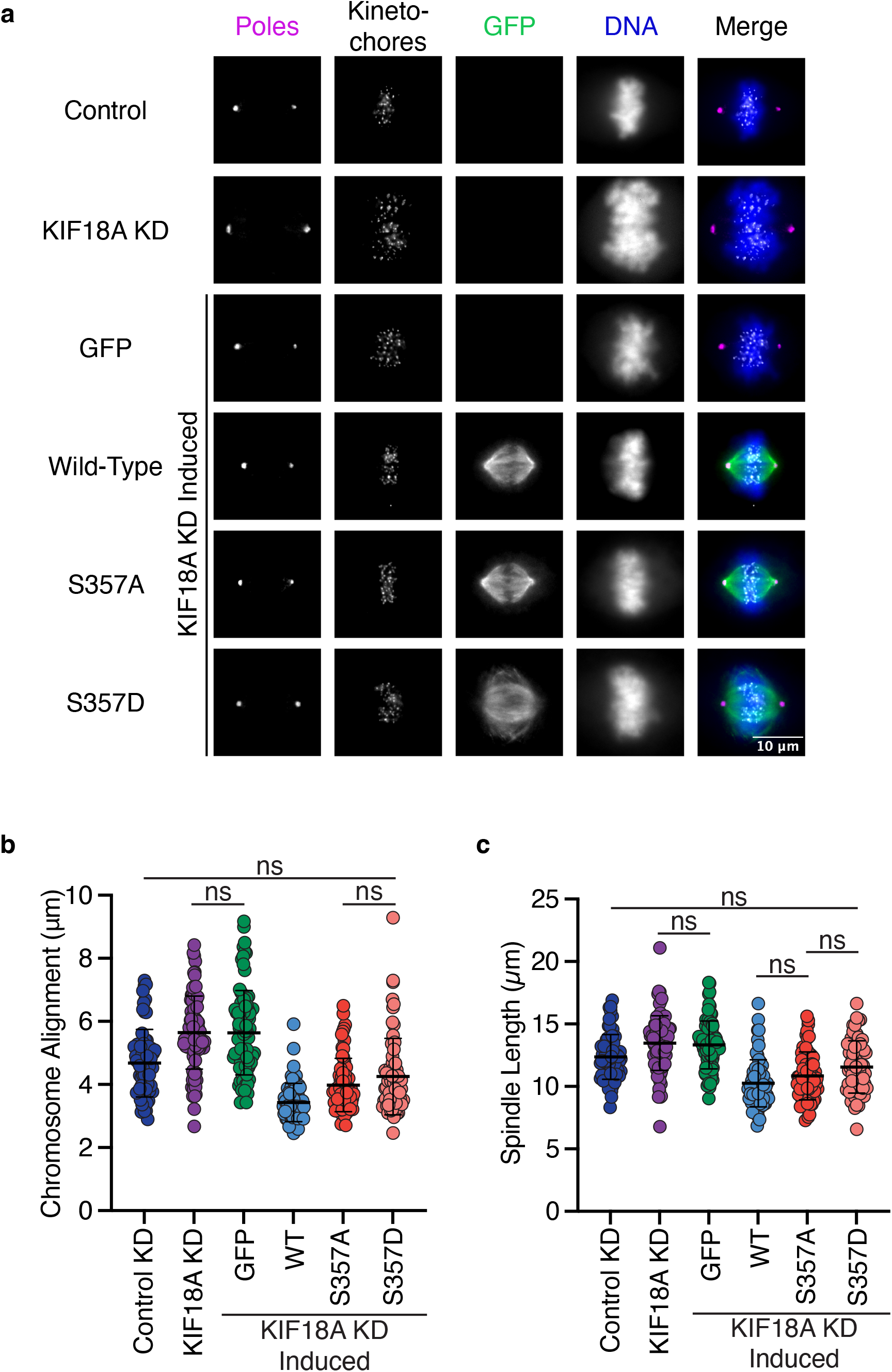
KIF18A S357D displays reduced chromosome alignment and spindle length control in RPE1 cells. **(a)** Representative immunofluorescence images of RPE1 cells. Cells were fixed approximately 24 hours after siRNA treatment to knockdown endogenous KIF18A and induction of GFP-KIF18A with doxycycline. Color indicates pseudo-color in merged image. Brightness/contrast levels for GFP are set to be equivalent, brightness/contrast levels for poles, kinetochore, and DNA are set differently to optimize visualization. Scale bar is 10 μm. KD: knockdown. **(b)** Quantification of chromosome alignment in RPE1 cells. Chromosome alignment was determined by measuring the distribution of kinetochores between spindle poles in a metaphase cell. This distribution was fit to a gaussian curve and the width of the distribution at half the maximum fluorescence intensity was recorded as the value for chromosome alignment. Solid horizontal line indicates mean, vertical lines indicate standard deviation. Each dot represents a single cell. Data were acquired from three experimental replicates and were analyzed using a one-way ANOVA with Tukey’s test for multiple comparisons. P value style: >0.05 (ns), if no significance is indicated result was significant (< 0.05). KD: knockdown, WT: wild-type. **(b)** Quantification of spindle length in RPE1 cells. Solid horizontal line indicates mean, vertical lines indicate standard deviation. Each dot represents a single cell. Data were acquired from three experimental replicates and analyzed using a one-way ANOVA with Tukey’s test for multiple comparisons. P value style: >0.05 (ns), if no significance is indicated result was significant (< 0.05). KD: knockdown, WT: wild-type.

## References

Atherton, J., J.J. Hummel, N. Olieric, J. Locke, A. Peña, S.S. Rosenfeld, M.O. Steinmetz, C.C. Hoogenraad, and C.A. Moores. 2020. The mechanism of kinesin inhibition by kinesin binding protein. Elife. 9:e61481. doi:10.7554/elife.61481.

Bachmann, A., and A. Straube. 2015. Kinesins in cell migration. Biochem Soc T. 43:79–83. doi:10.1042/bst20140280.

Barisic, M., R.S. e Sousa, S.K. Tripathy, M.M. Magiera, A.V. Zaytsev, A.L. Pereira, C. Janke, E.L. Grishchuk, and H. Maiato. 2015. Microtubule detyrosination guides chromosomes during mitosis. Science. 348:799–803. doi:10.1126/science.aaa5175.

Bickel, K.G., B.J. Mann, J.S. Waitzman, T.A. Poor, S.E. Rice, and P. Wadsworth. 2017. Src family kinase phosphorylation of the motor domain of the human kinesin-5, Eg5. Cytoskeleton. 74:317–330. doi:10.1002/cm.21380.

Brouhard, G.J., and L.M. Rice. 2018. Microtubule dynamics: an interplay of biochemistry and mechanics. Nat Rev Mol Cell Bio. 19:451–463. doi:10.1038/s41580-018-0009-y.

Brown, K.D., R.M.R. Coulson, T.J. Yen, and D.W. Cleveland. 1994. Cyclin-like Accumulation and Loss of the Putative Kinetochore Motor CENP-E Results from Coupling Continuous Synthesis with Specific Degradation at the End of Mitosis. J Cell Biol. 125:1303–1312. doi:10.1083/jcb.125.6.1303.

Cario, A., A. Savastano, N.B. Wood, Z. Liu, M.J. Previs, A.G. Hendricks, M. Zweckstetter, and C.L. Berger. 2022a. The pathogenic R5L mutation disrupts formation of Tau complexes on the microtubule by altering local N-terminal structure. Proc National Acad Sci. 119:e2114215119. doi:10.1073/pnas.2114215119.

Cario, A., S.P. Wickramasinghe, E. Rhoades, and C.L. Berger. 2022b. The N-terminal disease– associated R5L Tau mutation increases microtubule shrinkage rate due to disruption of microtubule-bound Tau patches. J Biol Chem. 298:102526. doi:10.1016/j.jbc.2022.102526.

Cohen-Sharir, Y., J.M. McFarland, M. Abdusamad, C. Marquis, S.V. Bernhard, M. Kazachkova, H. Tang, M.R. Ippolito, K. Laue, J. Zerbib, H.L.H. Malaby, A. Jones, L.-M. Stautmeister, I. Bockaj, R. Wardenaar, N. Lyons, A. Nagaraja, A.J. Bass, D.C.J. Spierings, F. Foijer, R. Beroukhim, S. Santaguida, T.R. Golub, J. Stumpff, Z. Storchová, and U. Ben-David. 2021. Aneuploidy renders cancer cells vulnerable to mitotic checkpoint inhibition. Nature. 590:486–491. doi:10.1038/s41586-020-03114-6.

Coy, D.L., W.O. Hancock, M. Wagenbach, and J. Howard. 1999. Kinesin’s tail domain is an inhibitory regulator of the motor domain. Nat Cell Biol. 1:288–292. doi:10.1038/13001.

Cross, R.A., and A. McAinsh. 2014. Prime movers: the mechanochemistry of mitotic kinesins. Nat Rev Mol Cell Bio. 15:257–271. doi:10.1038/nrm3768.

Czechanski, A., H. Kim, C. Byers, I. Greenstein, J. Stumpff, and L.G. Reinholdt. 2015. Kif18a is specifically required for mitotic progression during germ line development. Dev Biol. 402:253–262. doi:10.1016/j.ydbio.2015.03.011.

DeBerg, H.A., B.H. Blehm, J. Sheung, A.R. Thompson, C.S. Bookwalter, S.F. Torabi, T.A. Schroer, C.L. Berger, Y. Lu, K.M. Trybus, and P.R. Selvin. 2013. Motor Domain Phosphorylation Modulates Kinesin-1 Transport*. J Biol Chem. 288:32612–32621. doi:10.1074/jbc.m113.515510.

Deribe, Y.L., T. Pawson, and I. Dikic. 2010. Post-translational modifications in signal integration. Nat Struct Mol Biol. 17:666–672. doi:10.1038/nsmb.1842.

Dogan, M.Y., S. Can, F.B. Cleary, V. Purde, and A. Yildiz. 2015. Kinesin’s Front Head Is Gated by the Backward Orientation of Its Neck Linker. Cell Reports. 10:1967–1973. doi:10.1016/j.celrep.2015.02.061.

Du, Y., C.A. English, and R. Ohi. 2010. The Kinesin-8 Kif18A Dampens Microtubule Plus-End Dynamics. Curr Biol. 20:374–380. doi:10.1016/j.cub.2009.12.049.

Fink, J., N. Carpi, T. Betz, A. Bétard, M. Chebah, A. Azioune, M. Bornens, C. Sykes, L. Fetler, D. Cuvelier, and M. Piel. 2011. External forces control mitotic spindle positioning. Nat Cell Biol. 13:771–778. doi:10.1038/ncb2269.

Fonseca, C., and J. Stumpff. 2016. Quantification of Mitotic Chromosome Alignment. Methods in Molecular Biology. 253–262. doi:10.1007/978-1-4939-3542-0_16.

Fonseca, C.L., H.L.H. Malaby, L.A. Sepaniac, W. Martin, C. Byers, A. Czechanski, D. Messinger, M. Tang, R. Ohi, L.G. Reinholdt, and J. Stumpff. 2019. Mitotic chromosome alignment ensures mitotic fidelity by promoting interchromosomal compaction during anaphase. J Cell Biology. 218:1148–1163. doi:10.1083/jcb.201807228.

Funabiki, H., and A.W. Murray. 2000. The Xenopus Chromokinesin Xkid Is Essential for Metaphase Chromosome Alignment and Must Be Degraded to Allow Anaphase Chromosome Movement. Cell. 102:411–424. doi:10.1016/s0092-8674(00)00047-7.

Gardner, M., D. Odde, and K. Bloom. 2008. Kinesin-8 molecular motors: putting the brakes on chromosome oscillations. Trends in Cell Biology. doi:https://doi.org/10.1016/j.tcb.2008.05.003.

Garzon-Coral, C., H.A. Fantana, and J. Howard. 2016. A force-generating machinery maintains the spindle at the cell center during mitosis. Science. 352:1124–1127. doi:10.1126/science.aad9745.

Girão, H., and H. Maiato. 2019. Measurement of Microtubule Half-Life and Poleward Flux in the Mitotic Spindle by Photoactivation of Fluorescent Tubulin. Methods Mol Biology. 2101:235–246. doi:10.1007/978-1-0716-0219-5_15.

Goshima, G., R. Wollman, N. Stuurman, J.M. Scholey, and R.D. Vale. 2005. Length Control of the Metaphase Spindle. Curr Biol. 15:1979–1988. doi:10.1016/j.cub.2005.09.054.

Grill, S.W., and A.A. Hyman. 2005. Spindle Positioning by Cortical Pulling Forces. Dev Cell. 8:461–465. doi:10.1016/j.devcel.2005.03.014.

Hackney, D.D., J.D. Levitt, and J. Suhan. 1992. Kinesin Undergoes a 9 S to 6 S Conformational Transition*. J Biol Chem. 267:8696–8701. doi:https://doi.org/10.1016/S0021-9258(18)42499-4.

Hirokawa, N. 1998. Kinesin and Dynein Superfamily Proteins and the Mechanism of Organelle Transport. Science. 279:519–526. doi:10.1126/science.279.5350.519.

Hirokawa, N., Y. Noda, Y. Tanaka, and S. Niwa. 2009. Kinesin superfamily motor proteins and intracellular transport. Nat Rev Mol Cell Bio. 10:682–696. doi:10.1038/nrm2774.

Hirokawa, N., K.K. Pfister, H. Yorifuji, M.C. Wagner, S.T. Brady, and G.S. Bloom. 1989. Submolecular domains of bovine brain kinesin identified by electron microscopy and monoclonal antibody decoration. Cell. 56:867–878. doi:10.1016/0092-8674(89)90691-0.

Hoeprich, G.J., A.R. Thompson, D.P. McVicker, W.O. Hancock, and C.L. Berger. 2014. Kinesin’s Neck-Linker Determines its Ability to Navigate Obstacles on the Microtubule Surface. Biophys J. 106:1691–1700. doi:10.1016/j.bpj.2014.02.034.

Hornbeck, P.V., B. Zhang, B. Murray, J.M. Kornhauser, V. Latham, and E. Skrzypek. 2015. PhosphoSitePlus, 2014: mutations, PTMs and recalibrations. Nucleic Acids Res. 43:D512–D520. doi:10.1093/nar/gku1267.

Imami, K., N. Sugiyama, Y. Kyono, M. Tomita, and Y. Ishihama. 2008. Automated Phosphoproteome Analysis for Cultured Cancer Cells by Two-Dimensional NanoLC-MS Using a Calcined Titania/C18 Biphasic Column. Anal Sci. 24:161–166. doi:10.2116/analsci.24.161.

Jagrić, M., P. Risteski, J. Martinčić, A. Milas, and I.M. Tolić. 2021. Optogenetic control of PRC1 reveals its role in chromosome alignment on the spindle by overlap length-dependent forces. Elife. 10:e61170. doi:10.7554/elife.61170.

Janssen, L.M.E., T.V. Averink, V.A. Blomen, T.R. Brummelkamp, R.H. Medema, and J.A. Raaijmakers. 2018. Loss of Kif18A Results in Spindle Assembly Checkpoint Activation at Microtubule-Attached Kinetochores. Curr Biol. 28:2685–2696.e4. doi:10.1016/j.cub.2018.06.026.

Jumper, J., R. Evans, A. Pritzel, T. Green, M. Figurnov, O. Ronneberger, K. Tunyasuvunakool, R. Bates, A. Žídek, A. Potapenko, A. Bridgland, C. Meyer, S.A.A. Kohl, A.J. Ballard, A. Cowie, B. Romera-Paredes, S. Nikolov, R. Jain, J. Adler, T. Back, S. Petersen, D. Reiman, E. Clancy, M. Zielinski, M. Steinegger, M. Pacholska, T. Berghammer, S. Bodenstein, D. Silver, O. Vinyals, A.W. Senior, K. Kavukcuoglu, P. Kohli, and D. Hassabis. 2021. Highly accurate protein structure prediction with AlphaFold. Nature. 596:583–589. doi:10.1038/s41586-021-03819-2.

Karsenti, E., and I. Vernos. 2001. The Mitotic Spindle: A Self-Made Machine. Science. 294:543–547. doi:10.1126/science.1063488.

Kaul, N., V. Soppina, and K.J. Verhey. 2014. Effects of α-Tubulin K40 Acetylation and Detyrosination on Kinesin-1 Motility in a Purified System. Biophys J. 106:2636–2643. doi:10.1016/j.bpj.2014.05.008.

Kevenaar, J.T., S. Bianchi, M. van Spronsen, N. Olieric, J. Lipka, C.P. Frias, M. Mikhaylova, M. Harterink, N. Keijzer, P.S. Wulf, M. Hilbert, L.C. Kapitein, E. de Graaff, A. Ahkmanova, M.O. Steinmetz, and C.C. Hoogenraad. 2016. Kinesin-Binding Protein Controls Microtubule Dynamics and Cargo Trafficking by Regulating Kinesin Motor Activity. Curr Biol. 26:849– 861. doi:10.1016/j.cub.2016.01.048.

Khandelia, P., K. Yap, and E.V. Makeyev. 2011. Streamlined platform for short hairpin RNA interference and transgenesis in cultured mammalian cells. Proc National Acad Sci. 108:12799–12804. doi:10.1073/pnas.1103532108.

Koffa, M.D., C.M. Casanova, R. Santarella, T. Köcher, M. Wilm, and I.W. Mattaj. 2006. HURP Is Part of a Ran-Dependent Complex Involved in Spindle Formation. Curr Biol. 16:743–754. doi:10.1016/j.cub.2006.03.056.

Laan, L., N. Pavin, J. Husson, G. Romet-Lemonne, M. van Duijn, M.P. López, R.D. Vale, F. Jülicher, S.L. Reck-Peterson, and M. Dogterom. 2012. Cortical Dynein Controls Microtubule Dynamics to Generate Pulling Forces that Position Microtubule Asters. Cell. 148:502–514. doi:10.1016/j.cell.2012.01.007.

Malaby, H.L., D.V. Lessard, C.L. Berger, and J. Stumpff. 2019a. KIF18A’s neck linker permits navigation of microtubule-bound obstacles within the mitotic spindle. Life Sci Alliance. 2:e201800169. doi:10.26508/lsa.201800169.

Malaby, H.L.H., M.E. Dumas, R. Ohi, and J. Stumpff. 2019b. Kinesin-binding protein ensures accurate chromosome segregation by buffering KIF18A and KIF15. J Cell Biol. 218:jcb.201806195. doi:10.1083/jcb.201806195.

Marquis, C., C.L. Fonseca, K.A. Queen, L. Wood, S.E. Vandal, H.L.H. Malaby, J.E. Clayton, and J. Stumpff. 2021. Chromosomally unstable tumor cells specifically require KIF18A for proliferation. Nat Commun. 12:1213. doi:10.1038/s41467-021-21447-2.

Mayr, M.I., S. Hümmer, J. Bormann, T. Grüner, S. Adio, G. Woehlke, and T.U. Mayer. 2007. The Human Kinesin Kif18A Is a Motile Microtubule Depolymerase Essential for Chromosome Congression. Curr Biol. 17:488–498. doi:10.1016/j.cub.2007.02.036.

Muretta, J.M., B.J.N. Reddy, G. Scarabelli, A.F. Thompson, S. Jariwala, J. Major, M. Venere, J.N. Rich, B. Willard, D.D. Thomas, J. Stumpff, B.J. Grant, S.P. Gross, and S.S. Rosenfeld. 2018. A posttranslational modification of the mitotic kinesin Eg5 that enhances its mechanochemical coupling and alters its mitotic function. Proc National Acad Sci. 115:201718290. doi:10.1073/pnas.1718290115.

Nakata, T., and N. Hirokawa. 2003. Microtubules provide directional cues for polarized axonal transport through interaction with kinesin motor head. J Cell Biology. 162:1045–1055. doi:10.1083/jcb.200302175.

Nehlig, A., A. Molina, S. Rodrigues-Ferreira, S. Honoré, and C. Nahmias. 2017. Regulation of end-binding protein EB1 in the control of microtubule dynamics. Cell Mol Life Sci. 74:2381– 2393. doi:10.1007/s00018-017-2476-2.

Nishi, H., A. Shaytan, and A.R. Panchenko. 2014. Physicochemical mechanisms of protein regulation by phosphorylation. Frontiers Genetics. 5:270. doi:10.3389/fgene.2014.00270.

Olsen, J.V., B. Blagoev, F. Gnad, B. Macek, C. Kumar, P. Mortensen, and M. Mann. 2006. Global, In Vivo, and Site-Specific Phosphorylation Dynamics in Signaling Networks. Cell. 127:635–648. doi:10.1016/j.cell.2006.09.026.

Qian, L.-X., X. Cao, M.-Y. Du, C.-X. Ma, H.-M. Zhu, Y. Peng, X.-Y. Hu, X. He, and L. Yin. 2021. KIF18A knockdown reduces proliferation, migration, invasion and enhances radiosensitivity of esophageal cancer. Biochem Bioph Res Co. 557:192–198. doi:10.1016/j.bbrc.2021.04.020.

Quinton, R.J., A. DiDomizio, M.A. Vittoria, K. Kotýnková, C.J. Ticas, S. Patel, Y. Koga, J. Vakhshoorzadeh, N. Hermance, T.S. Kuroda, N. Parulekar, A.M. Taylor, A.L. Manning, J.D. Campbell, and N.J. Ganem. 2021. Whole-genome doubling confers unique genetic vulnerabilities on tumour cells. Nature. 590:492–497. doi:10.1038/s41586-020-03133-3.

Reed, N.A., D. Cai, T.L. Blasius, G.T. Jih, E. Meyhofer, J. Gaertig, and K.J. Verhey. 2006. Microtubule Acetylation Promotes Kinesin-1 Binding and Transport. Curr Biol. 16:2166– 2172. doi:10.1016/j.cub.2006.09.014.

Schindelin, J., I. Arganda-Carreras, E. Frise, V. Kaynig, M. Longair, T. Pietzsch, S. Preibisch, C. Rueden, S. Saalfeld, B. Schmid, J.-Y. Tinevez, D.J. White, V. Hartenstein, K. Eliceiri, P. Tomancak, and A. Cardona. 2012. Fiji: an open-source platform for biological-image analysis. Nat Methods. 9:676–682. doi:10.1038/nmeth.2019.

Schneider, C.A., W.S. Rasband, and K.W. Eliceiri. 2012. NIH Image to ImageJ: 25 years of image analysis. Nat Methods. 9:671–675. doi:10.1038/nmeth.2089.

Shastry, S., and W.O. Hancock. 2010. Neck Linker Length Determines the Degree of Processivity in Kinesin-1 and Kinesin-2 Motors. Curr Biol. 20:939–943. doi:10.1016/j.cub.2010.03.065.

Siahaan, V., R. Tan, T. Humhalova, L. Libusova, S.E. Lacey, T. Tan, M. Dacy, K.M. Ori- McKenney, R.J. McKenney, M. Braun, and Z. Lansky. 2022. Microtubule lattice spacing governs cohesive envelope formation of tau family proteins. Nat Chem Biol. 18:1224–1235. doi:10.1038/s41589-022-01096-2.

Silljé, H.H.W., S. Nagel, R. Körner, and E.A. Nigg. 2006. HURP Is a Ran-Importin β-Regulated Protein that Stabilizes Kinetochore Microtubules in the Vicinity of Chromosomes. Curr Biol. 16:731–742. doi:10.1016/j.cub.2006.02.070.

Sirajuddin, M., L.M. Rice, and R.D. Vale. 2014. Regulation of microtubule motors by tubulin isotypes and post-translational modifications. Nat Cell Biol. 16:335–344. doi:10.1038/ncb2920.

Solon, A.L., Z. Tan, K.L. Schutt, L. Jepsen, S.E. Haynes, A.I. Nesvizhskii, D. Sept, J. Stumpff, R. Ohi, and M.A. Cianfrocco. 2021. Kinesin-binding protein remodels the kinesin motor to prevent microtubule binding. Sci Adv. 7:eabj9812. doi:10.1126/sciadv.abj9812.

Stern, J.L., D.V. Lessard, G.J. Hoeprich, G.A. Morfini, and C.L. Berger. 2017. Phosphoregulation of Tau modulates inhibition of kinesin-1 motility. Mol Biol Cell. 28:1079–1087. doi:10.1091/mbc.e16-10-0728.

Stumpff, J., G. von Dassow, M. Wagenbach, C. Asbury, and L. Wordeman. 2008. The Kinesin-8 Motor Kif18A Suppresses Kinetochore Movements to Control Mitotic Chromosome Alignment. Dev Cell. 14:252–262. doi:10.1016/j.devcel.2007.11.014.

Stumpff, J., Y. Du, C.A. English, Z. Maliga, M. Wagenbach, C.L. Asbury, L. Wordeman, and R. Ohi. 2011. A Tethering Mechanism Controls the Processivity and Kinetochore-Microtubule Plus-End Enrichment of the Kinesin-8 Kif18A. Mol Cell. 43:764–775. doi:10.1016/j.molcel.2011.07.022.

Stumpff, J., M. Wagenbach, A. Franck, C.L. Asbury, and L. Wordeman. 2012. Kif18A and Chromokinesins Confine Centromere Movements via Microtubule Growth Suppression and Spatial Control of Kinetochore Tension. Dev Cell. 22:1017–1029. doi:10.1016/j.devcel.2012.02.013.

Sturgill, E.G., S.R. Norris, Y. Guo, and R. Ohi. 2016. Kinesin-5 inhibitor resistance is driven by kinesin-12Analysis of kinesin-5 inhibitor resistance. J Cell Biology. 213:213–227. doi:10.1083/jcb.201507036.

Vale, R.D., T.S. Reese, and M.P. Sheetz. 1985. Identification of a Novel Force-Generating Protein, Kinesin, Involved in Microtubule-Based Motility. Cell. 42:39–50. doi:10.1016/s0092-8674(85)80099-4.

Verhey, K.J., and J.W. Hammond. 2009. Traffic control: regulation of kinesin motors. Nat Rev Mol Cell Bio. 10:765–777. doi:10.1038/nrm2782.

Verhey, K.J., D.L. Lizotte, T. Abramson, L. Barenboim, B.J. Schnapp, and T.A. Rapoport. 1998. Light Chain– dependent Regulation of Kinesin’s Interaction with Microtubules. J Cell Biol. 143:1053–1066. doi:10.1083/jcb.143.4.1053.

Vicente, J.J., and L. Wordeman. 2015. Mitosis, microtubule dynamics and the evolution of kinesins. Exp Cell Res. 334:61–69. doi:10.1016/j.yexcr.2015.02.010.

Weaver, L.N., S.C. Ems-McClung, J.R. Stout, C. LeBlanc, S.L. Shaw, M.K. Gardner, and C.E. Walczak. 2011. Kif18A Uses a Microtubule Binding Site in the Tail for Plus-End Localization and Spindle Length Regulation. Curr Biol. 21:1500–1506. doi:10.1016/j.cub.2011.08.005.

Zhang, C., C. Zhu, H. Chen, L. Li, L. Guo, W. Jiang, and S.H. Lu. 2010. Kif18A is involved in human breast carcinogenesis. Carcinogenesis. 31:1676–1684. doi:10.1093/carcin/bgq134.

Zhu, C., J. Zhao, M. Bibikova, J.D. Leverson, E. Bossy-Wetzel, J.-B. Fan, R.T. Abraham, and W. Jiang. 2005. Functional Analysis of Human Microtubule-based Motor Proteins, the Kinesins and Dyneins, in Mitosis/Cytokinesis Using RNA Interference. Mol Biol Cell. 16:3187–3199. doi:10.1091/mbc.e05-02-0167.

